# Mutation Rate Evolution Drives Immune Escape In Mismatch Repair-Deficient Cancer

**DOI:** 10.1101/2022.03.06.482973

**Authors:** Hamzeh Kayhanian, Panagiotis Barmpoutis, Eszter Lakatos, William Cross, Giulio Caravagna, Luis Zapata, Kevin Litchfield, Christopher Steele, William Waddingham, Dominic Patel, Salvatore Milite, Chen Jin, Ann-Marie Baker, Christopher Ross, Daniel Alexander, Khurum Khan, Daniel Hochhauser, Marco Novelli, Benjamin Werner, Naomi Guppy, Josep Linares, Genomics England Research Consortium, Marjolijn J.L. Ligtenberg, Iris D. Nagtegaal, Andrea Sottoriva, Trevor Graham, Nischalan Pillay, Manuel Rodriguez-Justo, Kai-Keen Shiu, Marnix Jansen

## Abstract

Mutation rate optimisation drives evolution and immune evasion of bacteria and lentiviral strains, including HIV. Whether evolving cancer lineages similarly adapt mutation rates to increase tumour cell fitness is unknown. Here, by mapping the clonal topography of mismatch repair-deficient (MMRd) colorectal cancer, we show that genomic MMRd mutability co-evolves with neoantigen selection to drive intratumour diversification and immune escape. Mechanistically, we find that microsatellite instability modulates subclonal DNA repair by toggling two hypermutable mononucleotide homopolymer runs in the mismatch repair genes *MSH6* and *MSH3* (C8 and A8, respectively) through stochastic frameshift switching. Spontaneous mutation and reversion at these evolvability switches modulates subclonal mutation rate, mutation bias, and clonal HLA diversity during MMRd cancer evolution. Combined experimental and simulation studies demonstrate that subclonal immune selection favours incremental MMR mutations. MMRd cancers thus fuel intratumour heterogeneity by adapting subclonal mutation rate and mutation bias to immune selection, revealing a conserved co-evolutionary arms race between neoantigen selection and adaptive genomic mutability. Our work reveals layers of mutational complexity and microsatellite biology in MMRd cancer evolution previously hidden in bulk analyses.

## INTRODUCTION

Natural selection can tailor the mutation rate of a population by favouring allelic variants impairing DNA repair fidelity^1^. This hypermutator phenotype increases population diversity and therefore increases the likelihood that adaptive variants will be present within the population. Such hypermutator strategies are prevalent amongst bacterial strains to mediate adaptation to potentially lethal challenges^2, 3^. For example, chronic antibiotic management of patients with cystic fibrosis drives the selection of bacterial hypermutator lineages carrying antibiotic resistance genes^2^. In a similar fashion, HIV and other lentiviruses normally rely on the action of the virally-encoded Vif protein to counteract host antiviral APOBEC3G cytidine deaminase mutagenic activity^4^. However, naturally-occurring HIV variants with suboptimal anti-APOBEC3G activity drive a reservoir of hypermutated lineages which promote drug resistance and immune escape^5^. Although brief hypermutation can drive rapid adaptation, unchecked hypermutation is highly genotoxic and can negatively impact viability. Hypermutators are therefore generally counterselected following adaptation^3^, suggesting that fast evolving viruses and bacteria optimise mutation rates to adapt to local selection pressures^6^.

Bacterial species have evolved to balance the short-term adaptive advantages of hypermutation with its long-term mutagenic harms by restricting hypermutability to genomic loci most likely to mediate adaptation to selection constraints. So-called contingency loci contain hypermutable homopolymer sequences that stochastically fluctuate in repeat length^7, 8^. These phase-variable homopolymers act as evolutionary tuning knobs and provide an evolutionary advantage in bacteria of fast adaptation to new environments.

In human cells, post-replicative DNA mismatch repair (MMR) is performed by protein complexes consisting of MutS homolog 2 (MSH2) and MutS homolog 6 (MSH6), also known as MutS_α_, and MutL homolog 1 (MLH1) and PMS1 homolog 2 (PMS2), known as MutL_α_^9^. Alternatively, MSH2 can pair with MSH3 in a complex known as MutS_β_. MutS_α_ and MutS_β_ each function as DNA mismatch detection modules with partially overlapping specificities, whilst MutL_α_ (MLH1/PMS2) complexes execute mismatch repair. Notably, whilst the mutagenic impact of isolated MSH6 or MSH3 germline loss is relatively mild, combined MSH6/MSH3 inactivation in model systems drives a robust hypermutator phenotype^10^.

Loss of mismatch repair proficiency occurs in about 15% of colorectal cancers (CRCs) and is associated with the accumulation of single-nucleotide mismatches and frameshift variants due to insertion/deletion mutations in repetitive homopolymer sequences^11^. In the majority of cases this is due to *MLH1* hypermethylation^12^. The relentless accumulation of somatic variants renders growing mismatch repair-deficient (MMRd) tumours immunogenic and provokes extensive immuno-editing of evolving MMRd tumours^13^. Whilst many of the genetic targets associated with immune escape (such as HLA complex and B2M mutations) have been characterised^14^, the evolutionary trajectories and underlying processes that MMRd tumours take to navigate their immune selection landscape remain unknown, curtailing our ability to design strategies that predict or even thwart immune escape. Here we visualize the clonal architecture of evolving MMRd tumours to allow joint analysis of individual tumour subclones and the immune microenvironment at clonal resolution. We find that individual subclonal MMRd lineages harness hypermutable homopolymer sequences in *MSH6* and *MSH3* to adapt cellular mutation rate and mutation bias to subclonal immune selection. This strategy allows MMRd tumour subclones to engage in an evolutionary arms race with the evolving immune system and efficiently explore immune adaptation solutions, whilst minimising the deleterious impact of prolonged genomic hypermutation on cellular fitness.

## RESULTS

### *MSH6* and *MSH3* homopolymer frameshifts drive increased mutation burden

The loss of DNA mismatch repair in microsatellite unstable cancer drives tumour progression, but also provokes the relentless accumulation of neo-antigenic and deleterious mutations. We set out to investigate how growing MMRd cancers manage this balancing act between adaptive and deleterious mutations by screening for gene mutations that modulate tumour mutation burden in a large whole genome sequencing (WGS) dataset of 217 MMRd CRCs from the Genomics England 100,000 genomes project. Given that MMRd cancers predictably progress through the erosion of coding homopolymers in microsatellite instability (MSI) target genes such as *TGFBR2* (involved in TGFß-mediated growth inhibition) and *BAX* (involved in apoptosis regulation), we hypothesized the occurrence of secondary inactivating homopolymer mutations that correlate with mutation burden. To explore this, we carried out multiple linear regression analysis for the relationship between homopolymer frameshifts in MSI target genes and total mutation burden, controlling for patient age and tumour purity (details Supplementary Methods).

Homopolymer frameshift mutations in *MSH3* and *MSH6* revealed a strong positive correlation with mutation burden in this large MMRd CRC dataset (**Fig. 1A**). This result was surprising because, whilst frequent MSH6 and MSH3 homopolymer alterations have previously been described in large-scale compendium studies of MMRd cancers^15–17^, these homopolymer mutations had so far been disregarded because these were thought to be neutral passengers in the context of pre-existing (i.e truncal) MMR deficiency. As a control, we examined the relationship between homopolymer frameshift mutation of coding microsatellites in other frequently hit MMR gene targets in our multiple linear regression model, again controlling for patient age and tumour purity. This showed no correlation with mutation burden for any of these frequently hit targets, indicating that the positive correlation with mutation burden was specific to *MSH3* and *MSH6* homopolymer indels and not secondary to background gene mutation frequency (note legend **Fig. 1A** denoting frequency of homopolymer target gene mutation). Combined *MSH3* and *MSH6* frameshifts had an additive effect on mutation burden compared to either mutation alone (**Supplementary table 1**). Mutation of *MSH3* or *MSH6* increased mutation burden from a baseline estimate of 161,267 mutations by 88,038 and 63,675 mutations, respectively, whereas mutation of both was associated with an increase of 139,338 mutations. In these bulk sequencing data, frameshift mutation of *MSH3*, *MSH6,* or both was observed in 79% (172/217) of cases.

**Figure 1.**
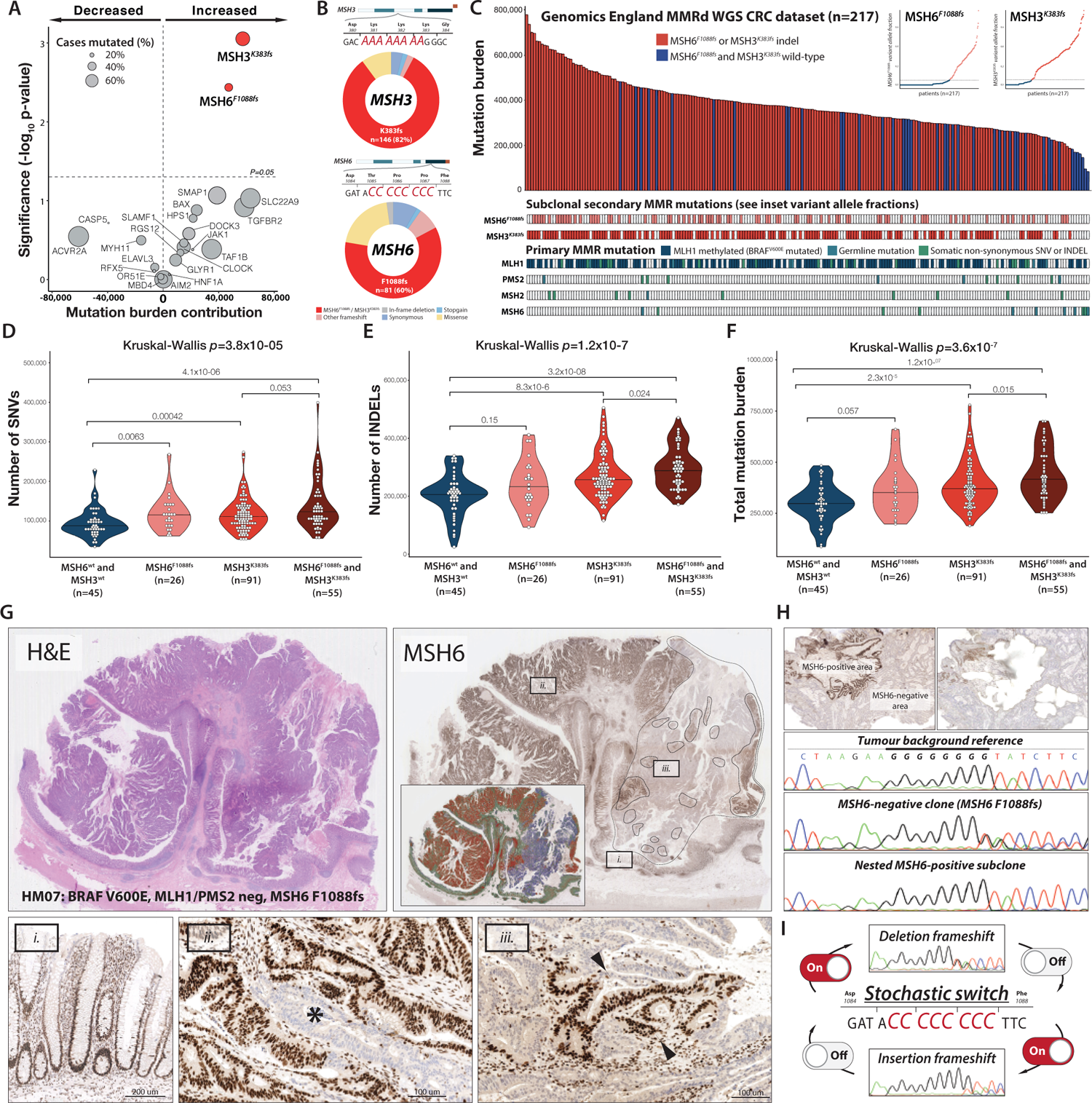
Subclonal *MSH6*^F1088fs^ and *MSH3^K383fs^* homopolymer frameshift mutations drive MMRd mutation burden. (A) Multiple linear regression showing relationship between homopolymer frameshifts in MSI target genes and increased or decreased mutation burden in the Genomics England MSI CRC cohort. (B) Pie charts showing mutation categories for *MSH3* (top) and *MSH6* (bottom). (C) Cases with *MSH6*^F1088fs^ and/or *MSH3^K383fs^* homopolymer frameshifts (in red) and cases without such mutations (in blue) in the Genomics England MSI CRC cohort (n=217) ranked by mutation burden. Clonal alterations in MMR genes MLH1, PMS2, MSH2 and MSH6, as well as subclonal *MSH6*^F1088fs^ and *MSH3^K383fs^* frameshift status is indicated in the panels below. Insets show *MSH6*^F1088fs^ and *MSH3^K383fs^* mutation variant allele fraction. (D-F) SNV, indel, and total mutation burden according to *MSH6*^F1088fs^ and *MSH3^K383fs^* mutation status. Median values are represented by horizontal black lines. (G) Example H&E and MSH6 immunohistochemistry for a case displaying subclonal loss of MSH6 expression. Inset shows segmentation for MSH6 labelling (in red) and MSH6 loss (in blue). Dashed areas indicate nested subclones showing restoration of MSH6 labelling. Detailed photomicrographs show normal crypts (i), focal loss of MSH6 labelling (ii), and nested subclone with restoration of MSH6 labelling (iii). (H) Laser capture microdissection (LCM) of tumour background, MSH6-negative and MSH6-positive nested area followed by Sanger sequencing of the MSH6 C8 coding microsatellite. (I) Sequential expansion and contraction of the hypermutable coding homopolymers in *MSH6* and *MSH3* unmasks these sites as cryptic stochastic switches in microsatellite unstable tumours.

*MSH6* and *MSH3* each contain a coding homopolymer tract (C8 and A8, respectively), which acts as a hypermutable site and is frequently mutated in MMRd tumours. The majority of *MSH6* and *MSH3* somatic mutations in the Genomics England MMRd CRC cohort occurred at these homopolymer sites (**Fig. 1B** and Supplementary table 2). As expected, these mutations are overwhelmingly subclonal events in this cohort as revealed by analysis of *MSH6* and *MSH3* homopolymer frameshift mutation variant allele frequencies (see inset **Fig. 1C**). As a further control, we compared the frequency of homopolymer frameshift mutations in these genes to the proportion of all exonic length 8 C:G or A:T homopolymers mutated across the cohort and found highly significant enrichment of homopolymer frameshift mutations in the *MSH6* C8 homopolymer (37.3% v 23.4%, *p*=1.9×10^-6^) and the MSH3 A8 homopolymer (67.3% v 9.9%; *p*<2.2×10^-16^).

Ranking our overall cohort by mutation burden illustrates the relationship between *MSH6* or *MSH3* homopolymer frameshift and overall tumour mutation burden (**Fig. 1C**). Next, stratifying by *MSH3* and/or *MSH6* homopolymer frameshift mutation status demonstrated a clear stepwise increase for SNVs, indels, and overall mutation burden. Notably, cases carrying combined *MSH6* and *MSH3* homopolymer mutations showed the highest mutation burden (burgundy), whilst cases carrying either *MSH6* (pink) or *MSH3* (red) mutation showed a comparable increase compared to tumours (blue) without *MSH6* or *MSH3* homopolymer mutation (**Fig. 1D-F**). As a further control we restricted our analysis to cases with confirmed truncal loss of MutL_α._ In MMRd tumours the presence of somatic *BRAF^V600E^* mutation associates strongly with *MLH1* promoter methylation and the presence of *BRAF^V600E^* is therefore an excellent predictor of *MLH1* hypermethylation in MMRd tumours^15, 18^. As expected, analysing cases with either *BRAF^V600E^* mutation or germline *MLH1* or *PMS2* pathogenic mutations (total n=135) further corroborated the relationship between MSH6^F1088fs^ / MSH3^K383fs^ and increased mutation burden (Fig. S1A-C). As further validation, we analysed TCGA whole exome sequencing data^15^. MSI cancers from the colorectal (n=48), uterine (n=67), stomach (n=63) and oesophageal (n=3) cohorts were identified and pooled. This analysis confirms the stepwise relationship for increased mutation and neoantigen burden in MSI tumours with *MSH6* and/or *MSH3* homopolymer frameshift mutations (Fig. S1D-I).

Together these data from two bulk sequencing cohorts suggest that secondary homopolymer frameshift mutations in *MSH3* and *MSH6* are a functional target of microsatellite instability and drive increased mutation burden. *MSH6* and *MSH3* homopolymer frameshift mutations are also frequently found in cell lines of MMRd tumours^19^. Indeed, previous in *vitro* work has indicated that *MSH6* homopolymer length varies between isogenic MMRd cell line isolates and fluctuates over time to drive spontaneous loss and restoration of MSH6 expression by moving in-and-out of reading frame^20^. Moreover, concomitant inactivation of MSH6 and MLH1 in isogenic cell lines increases cellular mutation rate compared to inactivation of MLH1 alone. These MSH6 homopolymer length fluctuations may thus provide a potential substrate for selection during tumour evolution, however this could not be evaluated in the *in vitro* context^20^.

Accurately determining the allelic status of *MSH6* and *MSH3* from bulk sequencing samples is complex due to the polymorphic nature of these homopolymers in clonal mixtures. However, immunohistochemistry (IHC) faithfully tracks MSH6 expression and is routinely used clinically to detect loss of MSH6 protein expression in MMRd tumours. To delineate the frequency of secondary MSH6 loss in a large clinical series, we prospectively investigated a series of 546 unselected CRCs using whole-slide MMR IHC (**Fig. S2A**). Of these cases, 88 (16%) were MMRd, of which 77 showed combined MLH1 and PMS2 loss, and 6 cases showed isolated PMS2 loss with intact MLH1. We found that 32 of these cases (37%) showed secondary MSH6 loss within the context of MLH1/PMS2 loss (**Fig. S2B,C**). Many of these tumours showed multiple geographically isolated MSH6-deficient subclones, which often varied substantially in size (**Fig. S2D**).

Remarkably, within MSH6-deficient clones we frequently found numerous nested subclones that had restored MSH6 expression (**Fig. 1G and Fig. S2D**). To verify that this nested patchwork faithfully reflected *MSH6* genotype, we carried out detailed laser capture microdissection (LCM) followed by Sanger sequencing of the background MSH6-proficient tumour, the MSH6-deficient lineage, and the MSH6-proficient nested subclone from each of three tumours. We indeed found sequential loss and restoration of the C8 homopolymer reading frame, suggesting that the *MSH6* homopolymer had dynamically expanded and contracted (C8>C9>C8) during tumour growth (**Fig. 1H**). Together, these data support the hypothesis that MMR homopolymer frameshifts act as a stochastic on/off switch, dynamically regulating MSH6 expression during MMRd tumour evolution, akin to bacterial contingency loci (**Fig. 1I**). We next set out to examine whether subclonal MMR homopolymer length fluctuations exacerbate intratumour heterogeneity and drive MMRd immune escape.

### MSH6 and MSH3 cooperatively shift substitution bias in MLH1/PMS2-deficient lineages

The spatially variegated pattern of *MSH6* labelling can be leveraged to interrogate the clonal landscape of MMRd tumours at monophyletic resolution. Using LCM we can directly assess the contribution of subclonal MSH6 and MSH3 homopolymer frameshift mutations to mutation burden within individual MSH6-proficient and MSH6-deficient tumour subclones and reconstruct highly resolved phylogenetic trees. We therefore collected multi-region whole exome sequencing (WXS) data after MSH6 IHC from 49 LCM regions (29 MSH6-proficient and 20 MSH6-deficient regions) from 22 MMRd tumours, including 11 MMRd tumours with heterogeneous MSH6 loss, and a control group of 11 MMRd MSH6-proficient cancers. In this dataset all cases came from stage II/III surgical resection specimens with no prior exposure to systemic therapy. All cases showed clonal loss of either MLH1 and PMS2, or PMS2 alone (from hereon collectively referred to as MLH1/PMS2 loss) and included patients with sporadic MLH1 inactivation and BRAF mutation as well as patients with germline MMR mutation. Detailed clinical and pathological characteristics are provided in **Supplementary table 3**.

Whole exome sequencing after LCM confirmed that tumour regions carrying frameshift mutations in either the *MSH6* or *MSH3* homopolymer had significantly increased SNV, indel, and overall mutation burden (**Fig. 2A-C**). As before, frameshift slippage of both *MSH6* and *MSH3* homopolymers had an additive effect. To account for the non-independence of multiple sampling per patient and the confounding impact of age and tumour purity on mutation burden, linear mixed-effect modelling was performed controlling for these variables. The effect of *MSH6^F1088fs^* and *MSH3^K383fs^* on total mutation burden remained significant after accounting for the random effect of individual variation between tumours and fixed effects of age and tumour purity (ANOVA *p*=0.0296, details **Supplementary table 4**). Large clinical cohorts have shown that tumour mutation burden in patients with *MSH2* and/or *MSH6* mutations is generally greater than in patients with *MLH1* and/or *PMS2* mutations^21^. Our data reveal that this relationship is recapitulated between individual tumour regions.

**Figure 2.**
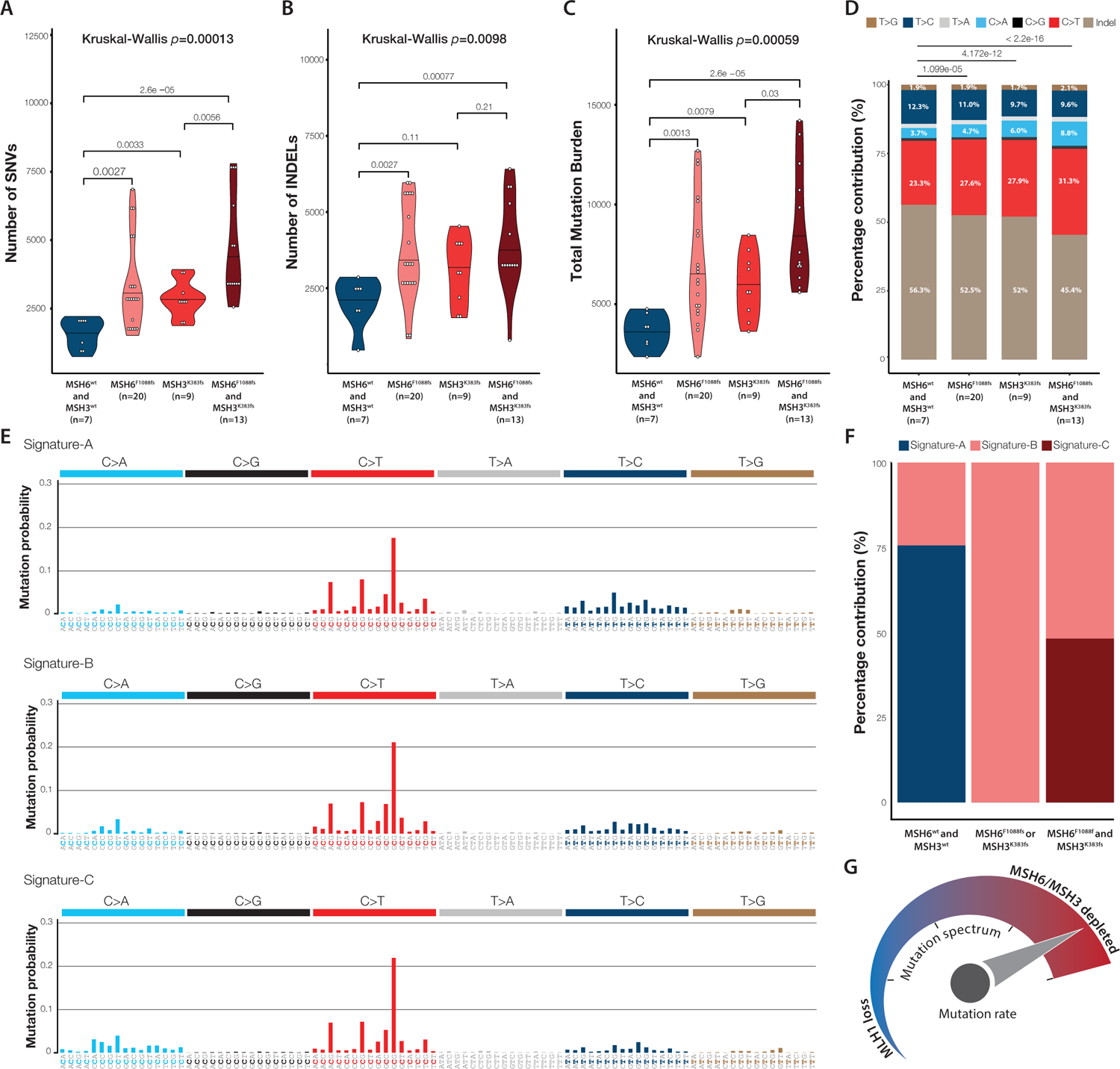
Subclonal *MSH6*^F1088fs^ and *MSH3^K383fs^* homopolymer frameshift mutations drive intratumour mutation burden and mutation bias heterogeneity. (A-C) SNV, indel, and total mutation burden in LCM samples according to *MSH6*^F1088fs^ and *MSH3^K383fs^* mutation status. (D) Mutation bias in samples according to *MSH6*^F1088fs^ and *MSH3^K383fs^* mutation status. (E) 96-Channel trinucleotide mutation spectra of de novo extracted mutation signatures. (F) Relative contribution of extracted mutation signatures according to *MSH6*^F1088fs^ and *MSH3^K383fs^* mutation status.(G) Subclonal *MSH6* and/or *MSH3* microsatellite frameshift mutations drive increased mutation rate and shift mutation bias towards *MSH2*-mutated phenotype.

Next, we compared mutation bias between regions. Data from a variety of model organisms ranging from *E.coli* to yeast and mice as well as from patients constitutively lacking mismatch repair have revealed that mutations of MutS_α_ or MutS_β_ drive a different mutation bias compared to mutations in MutL (MLH1/PMS2)^20, 22–24^. Specifically, whilst the mutation bias generated by MLH1/PMS2 loss is dominated by C>T and T>C transitions, MutS loss drives a decrease in T>C transitions in favour of C>T transitions and C>A transversions, along with a general increase in base substitutions over indels. Remarkably, analysis of the mutation spectrum in our microdissected samples recapitulated a clear stepwise decrease of T>C transitions and an increase in C>T transitions and C>A transversions in the presence of MSH6 and MSH3 frameshifts (*p*=1.099e-05, *p*=4.172e-12, and *p*<2.2e-16 for samples without secondary MSH6^F1088fs^ and MSH3^K383fs^ mutation compared to samples with secondary MSH6^F1088fs^, secondary MSH3^K383fs^, or both, respectively, **Fig. 2D**). In addition, whilst secondary MSH6/MHS3 frameshift mutations drove a combined increase in base substitutions as well as indels (**Fig. 2A-B**), the increase in base substitutions was markedly greater compared to indels (**Fig. 2D and Fig. S3**). These stepwise shifts in mutation bias recapitulate previous analyses comparing MutL and MutS_α/β_ mutation across model systems, and validate that incremental MMR mutations leave their indelible footprint on the genome by driving quantitative as well as qualitative mutation differences between tumour subclones.

We then carried out de novo signature deconvolution analysis on this sample cohort and retrieved three substitution signatures (here Signature-A, Signature-B, Signature-C, **Fig. 2E**) and three indel signatures (ID83-A, ID83-B and ID83-C, **Figure S4**). MLH1/PMS2 depletion without secondary MSH6/MSH3 loss is characterised by Signature-A exposure, whereas either *MSH6^F1088fs^* or *MSH3^K383fs^* mutation, or both, reveals Signature-B and Signature-C exposure, respectively (**Fig. 2F**). The shift in substitution bias towards an MutS_α/β_-dominated mutation bias with increasing MSH6 and MSH3 homopolymer frameshift mutation burden suggests that most MLH1/PMS2-deficient tumours are in fact a clonal mosaic of lineages with varying MMRd mutation biases, which is masked in bulk tumour analyses. The hypermutable homopolymers in *MSH6* and *MSH3* thus act as cryptic gear switches that are unmasked by microsatellite instability to fuel tumour heterogeneity (**Fig. 2G**).

### Secondary MMR mutations drive increased clonal diversity at a competing fitness cost

We set out to analyse the functional microenvironmental impact of secondary *MSH6* and *MSH3* homopolymer frameshift mutations. We reasoned that we could leverage our spatially deconvoluted LCM analysis by carrying out multiplex IHC labelling of key immune cell populations on serial sections to our LCM slides (**Fig. 3A**). We labelled for CD8, CD4, FoxP3, CD20, pan-cytokeratin, MSH6, and DAPI to trace immune cell infiltration and analysed 194 regions across 27 tumours. Accurate automated immune cell quantification in tumour cell regions is complicated by extensive overlapping of nuclei in standard tissue sections. In addition, normal immune cell populations express MSH6, necessitating strict separation of tumour cells from MSH6-positive immune cells. We therefore developed a dedicated fluorescence cell segmentation workflow (ORION) to accurately quantify immune cell infiltration both within and between MSH6-proficient and deficient tumour regions (**Fig. 3B-C, Fig. S5-6** and Methods for details).

**Figure 3.**
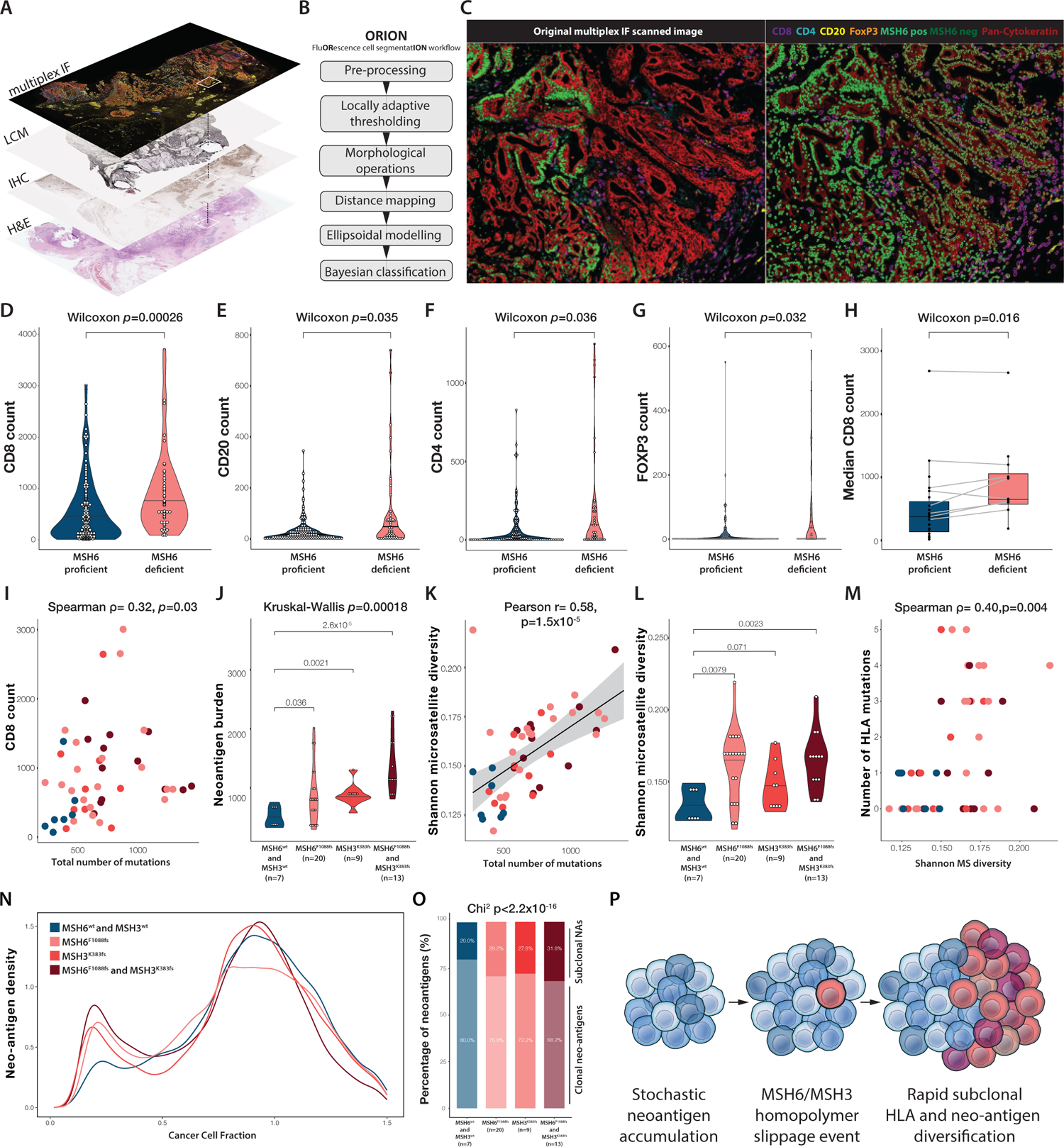
*MSH6*^F1088fs^ and *MSH3^K383fs^* homopolymer frameshift mutations accelerate clonal HLA diversity at the cost of increased neoantigen burden and immune cell infiltration. (A) Workflow for integrating MMRd clonal architecture, LCM sampling, and multiplex immunofluorescence experiments. (B) ORION (FluORescence cell segmentatION) workflow developed to investigate immune cell infiltration in MSH6-proficient and MSH6-deficient tumour subclones. (C) Example segmented multiplex immunofluorescence image showing the interface between MSH6-proficient and MSH6-deficient subclones. (D-G) Infiltration levels of CD8-pos, CD20-pos, CD4-pos and FOXP3-pos immune cells within 100μm radius of MSH6-proficient or MSH6-deficient tumour cells. (H) Median CD8 infiltration levels in MSH6-proficient and MSH6-deficient subclones of individual tumours. (I) CD8 count against total mutation burden according to *MSH6*^F1088fs^ and *MSH3^K383fs^* mutation status. Colour scheme as before. (J) Neoantigen burden in samples according to *MSH6*^F1088fs^ and *MSH3^K383fs^* mutation status. (K) Shannon population diversity of length 8 homopolymers against total mutation burden according to *MSH6*^F1088fs^ and *MSH3^K383fs^* mutation status. Colour scheme as before. (L) Shannon population diversity of length 8 homopolymers according to *MSH6*^F1088fs^ and *MSH3^K383fs^* mutation status. (M) Frequency of HLA class I mutations per sample according to *MSH6*^F1088fs^ and *MSH3^K383fs^* mutation status (Polysolver package). (N) Density plot showing fraction of neoantigens according to cancer cell fraction in samples grouped according to *MSH6*^F1088fs^ and *MSH3^K383fs^* status. (O) Percentage of clonal versus subclonal neoantigens in samples grouped according to *MSH6*^F1088fs^ and *MSH3^K383fs^* status. (P) In an MLH1/PMS2-deficient background, coding homopolymer slippage of MSH6 (C8) or MSH3 (A8) accelerates subclonal diversification. Colour scheme throughout as before.

Stratifying regions by MSH6 labelling revealed a clear increase in infiltrating CD8-positive cytotoxic T cells (**Fig. 3D**), CD20-positive B cells (**Fig. 3E**), CD4-positive T cells (**Fig. 3F**) and FoxP3-positive T-regulatory cells (**Fig. 3G**) in MSH6-deficient tumour regions. We then directly compared MSH6-proficient and deficient regions within individual tumours and again found increased CD8 infiltration in MSH6-deficient regions (**Fig. 3H**).

Next, we combined our clonally resolved WXS analyses with matched immune cell infiltration data and found that CD8 infiltration correlated with mutation burden (**Fig. 3I**). We then analysed neo-antigen burden again stratifying by secondary *MSH6^F1088fs^* and *MSH3^K383fs^* mutation status and observed a clear stepwise increase in neoantigen burden across these groups (**Fig. 3J**). These data corroborate our immune cell infiltration data comparing MSH6-proficient and deficient regions (**Fig. 3D-H**) and further suggest that subclonal neo-antigens drive intra-tumour differential immune cell infiltration in MMRd cancer.

We hypothesised that the drawbacks of increased neo-antigen burden and immune cell infiltration associated with *MSH6^F1088fs^* and *MSH3^K383fs^* might represent an evolutionary trade-off with the benefits conferred by greater subclonal genetic diversity, ultimately driving clonal selection and immune escape. We therefore queried the clonal diversity of our samples using lengths 6-11 exonic homopolymers as polymorphic lineage tags, further leveraging our clonal input LCM sampling strategy (**Fig. S7A-C**). We first examined the interaction between total mutation number and Shannon diversity and found a clear positive correlation (**Fig. 3K**). We then confirmed an increase in Shannon diversity across samples stratified by *MSH6^F1088fs^* and *MSH3^K383fs^* status (**Fig. 3L**). Finally, we analysed sample-specific HLA mutation status using Polysolver and found that broader clonal diversity correlated with a greater number of mutated HLA alleles (**Fig. 3M**). This result indicates that subclones marked by incremental MMR frameshift mutations reveal greater clonal diversity and mutation burden at key loci, which provides a substrate for immune selection and escape. To corroborate this analysis, we examined coding mutations in genes across the antigen presentation machinery (including B2M, JAK1, CITAA, and RFX5) and confirmed that subclones carrying *MSH6^F1088fs^* and *MSH3^K383fs^* indels show a greater number of antigen presentation machinery non-synonymous and indel mutations (**Fig. S8A-C**).

We then examined subclonal neoantigen complexity. Recent data from preclinical models^25^ as well as patient cohorts^26^ suggest that an increased number of subclonal neoantigens can drive early immune exhaustion and lack of checkpoint inhibitor treatment response. We find that subclones carrying either *MSH6^F1088fs^* or *MSH3^K383fs^* indels show a greater proportion of subclonal predicted neoantigens (**Fig. 3N-O**). Overall, these data indicate that evolving mismatch repair-deficient cancers elicit immune escape by balancing the evolutionary costs of adaptive hypermutability (increased deleterious genomic mutation load) against its fitness advantages (greater subclonal HLA and neoantigen diversity) (**Fig. 3P**).

### Adaptive hypermutability accelerates immune escape

Our data demonstrate that the hypermutable homopolymer sites in *MSH6* and *MSH3* act as cryptic genetic switches that are unmasked by microsatellite instability. The dynamic relationship between hypermutability at these key loci and immune adaptation in MMRd CRC is not a priori obvious. To understand the dynamic impact of mutation rate switching in a model wherein the ground truth is known, we extended our previous modelling work^13^ and created a model of a growing cancer that incorporates mutation rate switching through stochastic mutation and recovery of secondary MMR alleles. Briefly, the model captures early tumour growth (from 100 to 100,000 cells) using a stochastic birth-death process. Tumour cells can either die or proliferate according to their overall fitness, and accumulate new mutations during cell division, which may affect their fitness value (see model **Fig. 4A** and example scenarios **Fig. 4B**). Fitness is co-determined by the lineage-specific burden of stochastically accumulated neo-antigens and the prevailing immune selection *s*. Cells can either be in a basal MMRd hypermutated state (average 6 mutations gained per division) or ultra-hypermutated (average 120 mutations/division) state. The probability of switching between the two states is defined by the switch rate *β*, where *β =* 0 corresponds to mutation rates that remain constant and *β* = 0.01 represents frequent switching to or from the ultra-hypermutated state.

**Figure 4.**
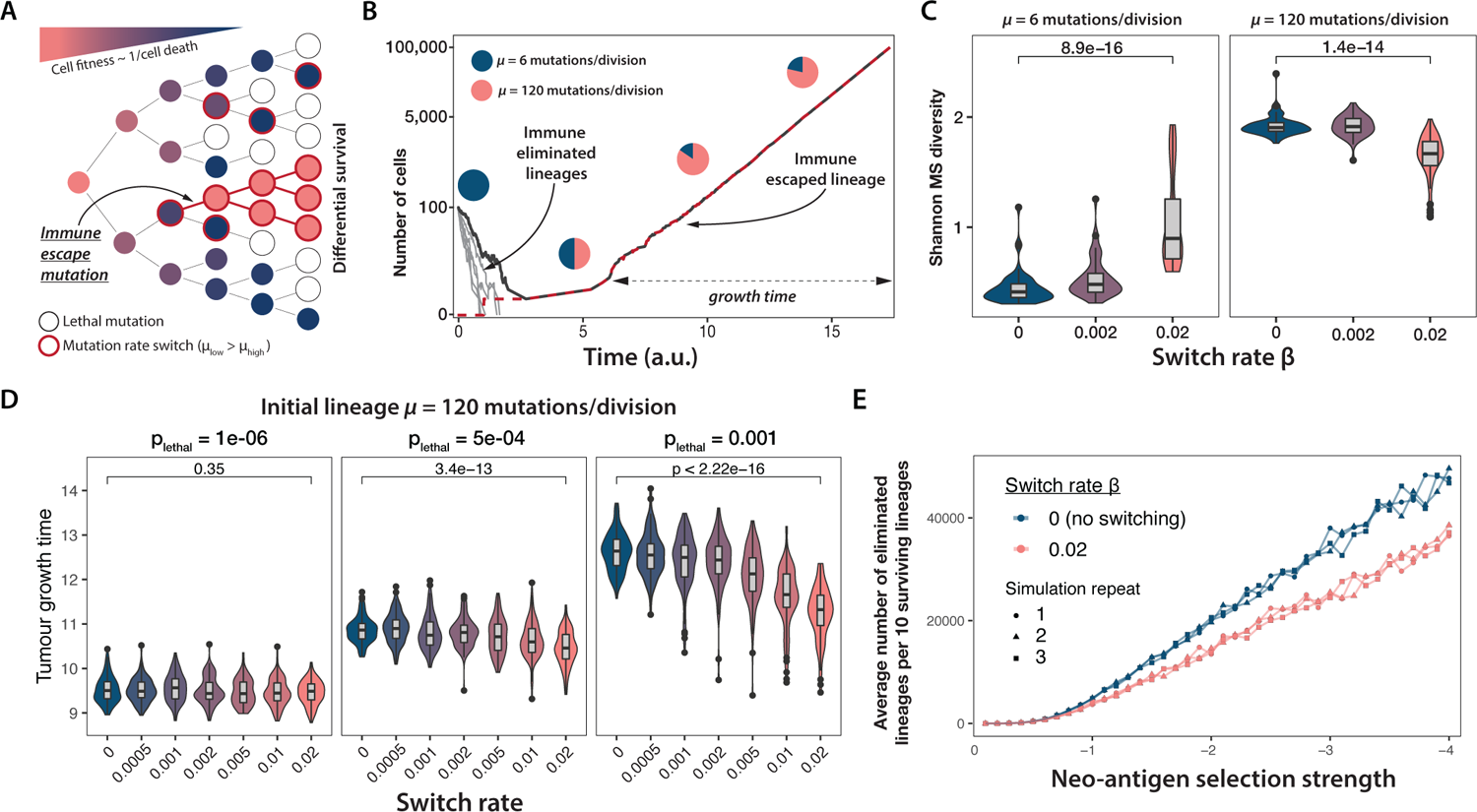
Mathematical model of the effect of stochastic mutation rate switching on tumour growth. (A) Schematic of the tumour growth model. The shade and outline colour of each circle (cell) represent that cell’s fitness and mutation rate, respectively. (B) Six simulated tumour growth trajectories. Five eliminated lineages are indicated in light grey, one surviving lineage in dark grey with the number of immune-escaped cells within the tumour shown in red (overlapping the dark grey curve). Pie-charts indicate the proportion of tumour cells with basal (blue) and higher (pink) mutation rate. (C) Shannon diversity of a microsatellite locus in 50 simulated tumours with varying starting mutation rate and switching rate. (D) Tumour growth time (in arbitrary units) between establishing immune escape and reaching detectable size (see model panel B), computed from 100 simulated tumours with starting mutation rate *m* = 120 mutations/division at increasing lethal mutation frequency (left to right) and mutation rate switching rate (x axis). The Wilcoxon-test statistic is reported on top of each panel in C and D. (E) Average (over 50 replicates) number of eliminated lineages per 10 surviving lineages as a function of selection strength, in tumours with no (=0, blue) and frequent (=0.02, pink) mutation rate switching. Three independent repeats of simulation and averaging are indicated by circle, triangles and squares.

First, we evaluated how mutation rate switching influences population diversity. We simulated the alterations of a single microsatellite locus throughout tumour growth and computed the overall Shannon diversity of this locus for each surviving detectable tumour. We found that higher mutation rates are associated with greater population diversity (**Fig. 4C**), corroborating observations within our clinical cohort (cf. **Fig. 3L**). Moreover, increased mutation rate switching either exacerbated or dampened population diversity depending on the mutation rate of the founding cell population (**Fig. 4C**).

To explore the impact of mutation rate switching on tumour evolution, we next examined tumour growth dynamics. We traced the number of lineages that were eliminated until 10 lineages had survived and repeated this analysis at different switch rates. As ultra-hypermutated cells have a higher probability of gaining an immune escape mutation, we hypothesised and confirmed that stochastic switching to a higher mutation rate decreases the rate of immunogenic lineage elimination. (**Fig. S9A**). Switching did not affect tumours with an ultra-hypermutated starting population (**Fig. S9B**), as these tumours already carried a significant proportion of fast mutating cells regardless of switching rate.

Following immune escape, tumours grow uninhibited by the antigenicity of rapidly accumulating mutations that characterise hyper-mutated tumours. However, other immune-independent deleterious mutations, represented in our model by lethal mutations, can still drastically decrease overall fitness. Since ultra-hypermutated cells acquire such disadvantageous mutations more frequently, we hypothesised that switching *back down* to a lower mutation rate is associated with a growth advantage for tumours initiated from these cells. We defined growth time as the time between complete immune escape and the tumour reaching clinically detectable size (**Fig. 4B**). We found that higher switching rates lead to a significantly faster tumour development and this effect becomes more pronounced as the likelihood of deleterious mutations increases (**Fig. 4D**). In tumours with a lower initial mutation rate, on the other hand, frequent switching leads to only small (although still significant) changes in growth time (**Fig. S9C**). Putting these results together, we examined the impact of immune selection and mutation rate switching on immune-mediated lineage extinction. We found that faster switching between mutation rates was associated with decreased lineage elimination which grew more prominent with increasing immune selection. Switching had no differential effect when immune selection was absent, as in this case lineages could reach detectable size without developing immune escape (**Fig. 4E**).

Altogether, these simulations indicate that mutation rate switching carries a survival advantage for hypermutated tumours. This interplay between opposing forces of deleterious mutation and immune extinction suggests that evolving MMR-deficient lineages undergo cycles of switching to and from an increased mutation rate, which maximises their survival and growth potential.

### Tumour phylogenies reveal frameshift reversion following immune escape

Next we wanted to explore the evolution of immune adaptation in individual MMRd tumours by harnessing the phylogenetic information contained within our multiregion WXS data. To this end we first established maximum parsimony phylogenetic trees from our multiregion SNV sequencing data (experimental workflow in **Fig. 5A**). We then evaluated immune escape alterations by joint inspection of tree topology and MSH6/MSH3 immunolabelling. As an internal control for immunolabelling, we plotted MSH6 and MSH3 homopolymer length traces from sequencing data using the MSIsensor package (**Fig. S11**). These plots show a wild-type reference allele (0 peak, in beige), representing stromal admixture and wild-type tumour alleles, and several additional contracted or expanded tumour alleles (in grey or red) depending on allelic drift within a sample (see example **Fig. 5A**). Phylogenetic analysis confirmed ongoing immune adaptation in growing MMRd tumours with many private immune escape mutations confined to tumour subclones (**Fig. 5B and S10**). As before, MSH6-deficient subclones carrying immune escape mutations frequently contained scattered nested subclones that had spontaneously restored MSH6 expression (see arrowheads **Fig. S10**), suggesting ongoing evolution of cellular mutability concurrent with subclonal immune adaptation. Our cohort represents a cross-sectional analysis of previously untreated MMRd tumours. These data suggest that untreated MMRd tumours typically explore multiple immune escape trajectories at diagnosis.

**Figure 5.**
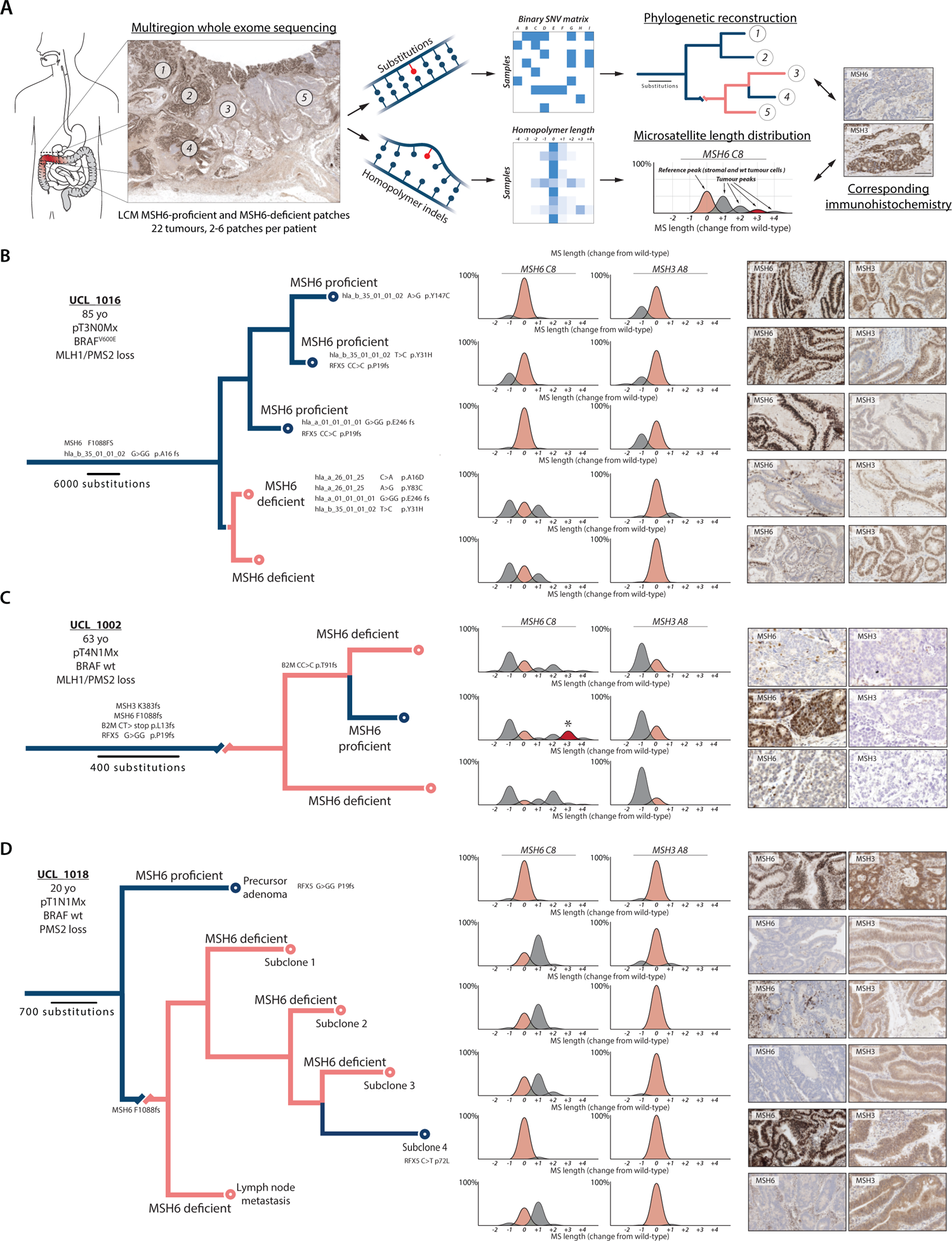
Phylogenetic trees reveal MSH6 homopolymer frameshift reversion events. (**A**) Cartoon showing workflow for phylogenetic reconstruction. Multi-region whole-exome sequencing data from MSH6-proficient and MSH6-deficient micro-dissected patches (22 tumours, between 2-6 patches per patient) are used to generate a binary SNV matrix to infer phylogenies using the maximum parsimony method (PAUP package). Scale bars indicate branch length and evolutionary distance expressed as substitution burden. The read length distribution of the MSH6 and MSH3 homopolymers are plotted separately (MSIsensor) and compared against the reference MSH6 and MSH3 immunohistochemistry of the input sample. (B-D) Maximum parsimony phylogenetic trees were reconstructed using SNV mutation data. Branch length is proportional to the number of mutations. Inset shows clinicopathological characteristics. Branches coloured according to MSH6 IHC labelling, blue indicating MSH6-proficient and pink indicating MSH6-deficient lineages. High-power photomicrographs show MSH6, MSH3 and MSH2 labelling as indicated. Trees are labelled with pertinent immune escape mutations. Microsatellite length distribution shows MSIsensor output where peak height is proportional to allelic frequency with beige indicating wild-type C8 or A8 length, dark grey indicating expanded or contracted alleles, and red indicating +3 frameshift.

We then evaluated temporal evolution of secondary MMR mutations from these phylogenetic trees. This analysis provided direct support for MSH6 homopolymer frameshift switching during tumour evolution. **Figure 5C** depicts a phylogeny consisting of three terminal branches, two of which derive from MSH6-deficient patches, while the third derives from a MSH6-proficient patch. All patches revealed loss of MSH3 labelling. MSH2 labelling was lost in regions that had lost both MSH6 and MSH3, while it was restored in the MSH6-proficient patch, indicating that simultaneous loss of MSH6 and MSH3 expression leads to complete loss of MSH2 labelling (**Fig. S12**). Phylogenetic ordering revealed that the MSH6-proficient lineage derived from the MSH6-deficient clade, providing formal proof of frameshift restoration. Analysis of the MSH6 homopolymer length distribution showed that the MSH6-proficient lineage carried a +3 insertion (see asterisk **Fig. 5C**), indicating the *MSH6* homopolymer had undergone progressive nucleotide insertion until the reading frame was restored after which the lineage clonally expanded. Notably, this reverter lineage carried a *B2M* mutation, providing a potential explanation for its selection and clonal expansion. In a second example (**Fig. 5D and S12**) phylogenetic ordering again revealed an MSH6-proficient branch that derived from an MSH6-deficient clade. The reverted clade in this case carried a non-synonymous C>T mutation in the HLA class II regulatory protein RFX5. Taken together, these phylogenetic data corroborate our model simulations and show that MMRd tumours exploit the inherent hypermutability of the *MSH6* and *MSH3* homopolymer coding sequences to drive diverse immune escape solutions. Multiple divergent paths are found within individual MMRd tumours reflecting the stochastic nature of subclonal neo-antigen generation and immune selection.

### MMRd tumours are a clonal mosaic of mutation rates

Hypermutator variants in lower organisms are well known to drive accelerated adaptation to ecological constraints at a competing fitness cost^8, 27^. These mutator lineages change the fitness distribution of their descendants and thus hitchhike with the adaptive variants they generate, rather than being directly selected^2, 3^. *MSH6^F1088fs^* and *MSH3^K383fs^* drive increased clonal HLA diversity providing a potential mechanism for clonal selection, but whether immune selection favours secondary MutS mutations is unclear. We sought to test this directly using immune dNdS analysis. Immune dNdS measures the ratio of nonsynonymous to synonymous mutations at genomic loci that are exposed to the immune system (Immune ON) compared to neutral expectation. Here we calculated dNdS across genomic regions that bind to HLA-A0201, the most common HLA class I allele in the Caucasian population, to compare patients across our cohort. Immune ON dNdS is compared to dNdS values in regions outside of those exposed to the immune system (Immune OFF) as an internal control (**Supplementary table 7**). We first examined tumour regions without *MSH6^F1088fs^* or *MSH3^K383fs^* and found no significant difference in nonsynonymous mutation accumulation between regions exposed and not exposed to the immune system (**Fig. 6A**). By contrast, regions with *MSH6^F1088fs^* and/or *MSH3^K383fs^* showed a highly significant difference when comparing immune ON and OFF dNdS (**Fig. 6A**). Enrichment of nonsynonymous mutations in HLA-A0201-binding regions in lineages with *MSH6^F1088fs^* and/or *MSH3^K383fs^* supports selection of these variants through linked immune escape variants.

**Figure 6.**
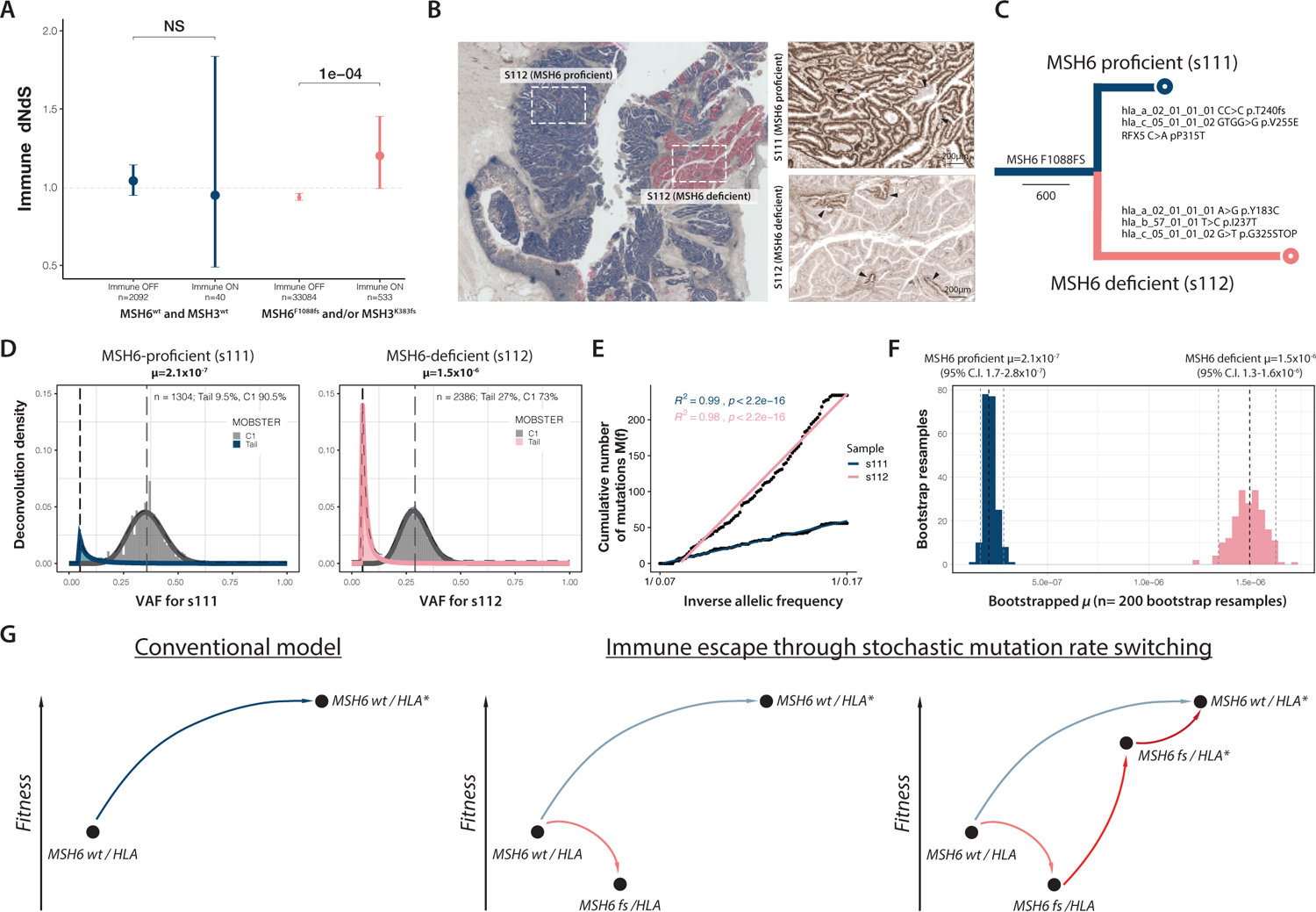
Clonal deconvolution quantifies elevated mutation rate in MSH6-deficient subclones. (A) Immune dNdS scores according to *MSH6*^F1088fs^ and *MSH3^K383fs^* mutation status. (B) Overview case UCL_1014 with LCM samples as indicated. Arrowheads indicate minute reverter clones. (C) Phylogenetic tree for tumour UCL_1014 annotated with HLA mutations identified in samples s111 and s112, respectively. (D) MOBSTER subclonal deconvolution from diploid variants detected in LCM samples s111 (left) and s112 (right) from patient UCL_1014 shows a monoclonal population with a tail of neutral mutations. (E) Cumulative frequency distribution of subclonal tail mutations in s111 and s112. The point estimate of the normalised mutation rate μ in panel F is estimated from the slope of the cumulative frequency distribution. (F) Bootstrapped percentile CI for the point estimate of the mutation rates in panel E (G) Model illustrating evolutionary trajectories to immune escape in MMRd cancer. Secondary *MSH6*^F1088fs^ and *MSH3^K383fs^* mutations initially decrease fitness (increased neoantigen burden and immune cell infiltration), but increase allelic diversity thereby decreasing the waiting time to immune escape (indicated by HLA*). Following immune escape, normal reading frame is restored (MSH6^wt^).

As a final test we set out to derive clone-specific *in vivo* mutation rates comparing MSH6-proficient and MSH6-deficient regions in individual tumours. In order to retrieve *in vivo* mutation rate estimates from single timepoint sequencing data, we developed a pipeline around MOBSTER, a recently published computational method that performs tumour subclonal deconvolution by integrating population genetics and machine learning^28^. This method retrieves a mutation rate estimate from the tail of neutral mutations within the allele frequency spectrum of a given sample. Specifically, we applied a population genetics approach to estimate sample-specific mutation rates *μ* from the fit of neutral tails, which we normalised for the portion of the diploid genome that had undergone WXS sequencing (see Methods for details). Comparison of mutation rate values between samples of patient UCL-1014 (**Fig. 6B,C**) shows that *μ* = 2.11^-7^ for the MSH6-proficient sample, whilst *μ* =1.51^-6^ for the MH6-deficient sample (**Fig. 6D-F**). The confidence intervals (bootstrapped percentile from 200 resamples, see Methods) for both estimates were non-overlapping with a difference of over one order of magnitude between *μ* point estimates (**Fig. 6F**). These results further corroborate the mutator dynamics from the simulated branching process described in Figure 4 and provide direct evidence that MMRd tumours are a mosaic of varying mutation rates.

## DISCUSSION

The distribution of somatic mutations across cancer genomes is highly heterogeneous and influenced by a complex interaction of factors including replication timing, chromatin compaction and three-dimensional genome organisation^29, 30^. The mismatch repair system plays a pivotal role in this regard^31, 32^. Recent mutational analyses of normal tissues from patients constitutively lacking mismatch repair have shown that, compared to MLH1 loss, MSH6 loss markedly increases substitution rates and disproportionately drives mutation accumulation in early-replicating protein-coding regions^24^, presumably because of the direct engagement of MSH6 with the H3K36me3 chromatin mark enriched on exons^33^. Our data corroborate these analyses showing that individual members of the mismatch repair pathway promulgate distinct mutation biases, which can become a substrate for subclonal selection during cancer evolution.

Here we visualize the clonal architecture of evolving MMRd tumours to allow joint analysis of individual tumour subclones and the immune microenvironment at clonal resolution. The subclonal evolution of MMRd tumours is characterized by the relentless accumulation of neoantigenic and deleterious variants, which drives a complex mosaic of divergent subclones carrying distinct neo-antigen profiles^13^. We find that subclonal MMRd lineages adapt to immune selection by repurposing hypermutable mononucleotide repeats in the *MSH6* and *MSH3* coding sequences as bistable switches to drive stochastic loss of expression at these loci. Our bulk sample and laser capture dissection data show that incremental MMR mutations increase cellular mutation rate, redirect mutation bias, and increase clonal diversity. This expands accessible genotype space and increases population diversity, allowing natural selection to pick immune-adapted variants (Fig. 6G).

This strategy is not without risk, however, because the majority of novel variants with phenotypic effects are deleterious^34^. The combined impact of deleterious mutation and accelerated drift at key immune adaptation loci might thus quickly negate prior fitness gains and prolonged hypermutation is therefore costly. Previous work in lower organisms shows that unmitigated hypermutation can drive clonal extinction through the irreversible accumulation of deleterious variants in a process known as Muller’s ratchet^35^. Remarkably, we find that expanding lineages carrying biallelic homopolymer frameshift mutations stochastically produce nested subclones marked by restored MSH6 protein expression. This corroborates previous *in vitro* work which showed that the *MSH6* homopolymer drifts over time in MLH1-deficient MMRd colorectal cancer cell lines redirecting mutation rate and bias^20^. This bet-hedging strategy allows parent lineages to continue exploring the local fitness landscape, whilst daughter lineages expand at lower mutation rates, hedging against the mutagenic harms associated with unmitigated hypermutation and acting as a reservoir to retain prior fitness gains. Comparable strategies driving lineage survival through punctuated burst of accelerated mutability have been extensively characterized in lower organisms and commonly target coding homopolymers in mismatch repair homologues^8, 27^.

Overall, our data reveal that evolving MMRd CRCs engage in a strategic evolutionary arms race between adaptive hypermutability and immune selection. This ancient survival strategy exploits the recurrently hit coding homopolymers of *MSH6* and *MSH3* as genomic evolvability switches. A mutator strategy exacerbates intratumour heterogeneity and increases subclonal neo-antigen burden. Recent preclinical studies indicate that increased subclonal complexity and subclonal neoantigen burdens blunt immune responses to checkpoint inhibitors in MMRd animal models due to immune exhaustion^25^. In line with this, we have recently found that low neoantigen clonality predicts lack of response to checkpoint inhibitors in MMRd patients^26^. Together this suggests that increased HLA diversity and subclonal neoantigen complexity jointly shape immune evasion in MMRd cancer.

Drug selection experiments in microsatellite-stable colorectal cancer have provided evidence for adaptive mutability in response to treatment^36^. Our work adds to this evidence base and reveals that adaptive mutability drives lineage survival during MMRd cancer evolution prior to clinical intervention. This evolutionary strategy – balancing clonal diversity against mutation load – matches hard-wired survival strategies of lower organisms and viruses driving resistance evolution^6, 37^. Understanding the evolutionary pathways to immune escape in MMRd tumours may allow us to map and forecast individual responses to immune checkpoint inhibition^38^.

## Acknowledgements

This research was made possible through access to the data and findings generated by the 100,000 Genomes Project. The 100,000 Genomes Project is managed by Genomics England Limited (a wholly owned company of the Department of Health and Social Care). The 100,000 Genomes Project is funded by the National Institute for Health Research and NHS England. The Wellcome Trust, Cancer Research UK and the Medical Research Council have also funded research infrastructure. The 100,000 Genomes Project uses data provided by patients and collected by the National Health Service as part of their care and support. We thank all patients who contributed samples to this study. Support was provided to the Jansen lab by the National Institute for Health Research, the University College London Hospitals Biomedical Research Centre, and the Cancer Research UK University College London Experimental Cancer Medicine Centre. We also acknowledge the pathology, genomic, and microscopy Translational Technology Platforms (TTP) at UCL Cancer Institute.

## Author contributions

MJ and HK conceived the project and designed the experimental strategy. HK performed the wet lab experiments. KL performed variant calling of sequencing data. HK performed downstream bioinformatic analysis. PB performed image analysis of multiplex immunofluorescence data. EL performed mathematical modelling experiments. GC performed MOBSTER clonal deconvolution and mutation rate analysis. CS provided advice on statistical analysis. MJ, HK, GC, EL, WC and KKS contributed to writing the manuscript. All authors edited and approved the final manuscript.

## Supplementary methods

### UCL Colorectal Cancer Cohort

All samples were processed with patient consent, according to protocols approved by the UCL/UCLH Biobank of health and disease ethical review committee (Project Reference Number NC21.18). The biobank was searched to identify MMRd colorectal cancers diagnosed between 2014 to 2018. Of 546 cancers tested by immunohistochemistry, 88 (16%) showed mismatch repair protein loss. Available FFPE tumour blocks were retrieved and sections cut to perform MSH6 immunohistochemistry (IHC) using an established protocol. Antibody details and immunohistochemistry conditions are provided in Table S3.

After assessing for tissue quality, eleven (n=11/40, 28%) tumours had subclonal loss of MSH6 in at least one tumour block. A stage and age-matched cohort of 11 MLH1/PMS2 MMR-D tumours without immunohistochemical loss of MSH6 was also selected as the comparison group. For each tumour, a corresponding normal block from the resection margin was also retrieved. MSH6 labelled slides for each tumour were scanned using a slide scanner (Hamamatsu NanoZoomer).

### Laser Capture Microdissection (LCM)

Tumours with MSH6 deficient subclones (n=11) and those in the MSH6 proficient comparison group (n=11) were taken forward for laser capture microdissection (LCM). Multi-region samples from more than one tumour block were taken where available. Since IHC labelling can affect DNA yield, a bespoke protocol was developed so that adjacent IHC labelled sections were used to guide microdissection of thicker sections on LCM membrane slides. Each tumour block was serially sectioned as follows: one 3um thick section on to a glass slide, five 10um thick sections onto poly-ethylene naphtholate (PEN) membrane slides (Zeiss Ag) and one 3um thick section on to a glass slide. The 3um thick sections underwent IHC against MSH6 and were used to guide microdissection of the thicker sections in between. Membrane slides were pre-treated with 0.01% poly-l-lysine to improve tissue adherence. Mounted sections were baked in an oven at 50°C for 4 hours. The 10um thick sections were deparaffinized and stained with haematoxylin as follows: xylene (10 minutes, two changes), 100% ethanol (1 minute, two changes), 90% ethanol (1 minute, one change), rinse in deionized water, Gill’s haematoxylin (1 minute, one change), rinse gently in running water, 90% ethanol (1 minute, two changes), 100% ethanol (1 minute, two changes), xylene (1 minute, two changes). LCM was performed using the Palm Microbeam microscope (Zeiss Ag). Selected MSH6 deficient and proficient tumour regions approximately 2-3mm^2^ in area were individually microdissected and collected in 500uL AdhesiveCap tubes (Zeiss Ag). Tissue originating from the same location was pooled across serial sections and processed as one sample. Tissue from the resection margin normal mucosa was also microdissected and used as the germline sample. Each microdissected region was allocated a unique sample number and the microdissected site recorded on corresponding scanned slides for future reference.

### Immunohistochemistry

Immunohistochemistry was performed using the Leica Bond autostainer (Leica Biosystems). Antibody details and conditions are provided in Supplementary table 5.

### DNA extraction

6ul of proteinase K and 200ul of lysis buffer (Perkin Elmer Inc.) was added to micro-dissected tissue samples and incubated overnight at 56°C followed by 1 hour at 70°C to reverse formaldehyde crosslinks. DNA extraction was completed using the Chemagic Prepito automated instrument (Perkin Elmer Inc) which uses a magnetic particle separation technique. Extracted DNA was quantified using a Qubit fluorometer (Thermo Fisher) as per the manufacturer’s instructions.

### Sanger Sequencing

Validation of frameshift mutation in the C8 coding microsatellite within *MSH6* was performed by PCR followed by BigDye terminator Sanger sequencing. Oligonucleotides (Forward primer TTTTAACAGATGTTTTACTGTGC and Reverse primer TCATTAGGAATAAAATCATCTCC), Q5 polymerase mastermix (New England Biolabs) and 10ng of DNA were used in PCR reactions. PCR conditions were 35 cycles of denaturation at 95°C for 30 seconds, followed by primer annealing at 60°C for 1 minute, followed by extension at 72°C for 30 seconds.

### Sample preparation for Whole Exome Sequencing

Acoustic fragmentation of DNA was performed using the Covaris E220 device. 125ng of sample DNA was inserted into snap-cap microtubes (Covaris) at a total volume of 50uL. The Covaris device was used as per the manufacturer’s guidelines. The following settings were used: Duty factor=10%, peak incident power (W)=175, cycles per burst=200, time (seconds)=300. Fragmented DNA samples were transferred to 1.5ml Eppendorf tubes.

### FFPE repair

FFPE repair was performed to minimize impact of artefactual lesions due to formalin fixation (1) using a validated kit (M6630L, New England Biolabs) and following the manufacturer’s protocol. Briefly 48ul of fragmented sample DNA was mixed with 3.5ul of FFPE DNA repair buffer, 3.5ul of end-prep buffer and 2ul of FFPE DNA repair mix and the mixture incubated at 20°C for 30 minutes.

### Library preparation

Library preparation was performed using the NebNext Ultra II kit (New England Biolabs) as per the manufacturer’s protocol. Briefly, following end repair and A-tailing, adapter ligation was performed by adding 30ul of ligation master mix, 1ul of ligation enhancer and 2.5ul of sequencing adapters and the reaction incubated for 15 minutes at 20°C. Adapters were diluted 10x as per manufacturer’s guidance. Magnetic bead clean-up of adapter ligated libraries was performed by adding 87ul (0.9x) of Ampure XP beads (Beckman Coulter) followed by ethanol washes and elution in 17ul of 10mM Tris-HCl. Next 15ul of adapter ligated library was amplified with 10 cycles of PCR by adding 25uL Q5 master mix and 10ul of indexing primers. For sample indexing NEBNext Multiplex Oligos (#E7335) were used and the indexing primer used recorded for each sample. Library fragment size was analysed with a Tapestation device (Agilent Technologies) using the high sensitivity screentape and also quantified using a Qubit fluorometer (Thermo Fisher).

### Exome capture

Exome capture was performed following the manufacturer’s protocol in the SeqCap EZ kit (Roche Sequencing Solutions). 250ng of library samples from the previous step were pooled in groups of four to give a total mass of 1ug. The multiplexed library pool was hybridized with SeqCap EZ Prime Exome probes for 16 hours at 47°C. Following hybridization, unbound probes were washed away and the hybridized DNA was amplified with 14 cycles of PCR followed by 1x Ampure XP bead clean up and eluted in 33ul of 0.1x TE solution. The final captured amplified library was quantified by qPCR using the NEBNext Library Quant kit for Illumina (New England Biolabs).

### Next Generation Sequencing

Samples were diluted to 2nM and sequenced in batches of 12 samples on the NovaSeq instrument (Illumina) using an S1 flowcell with 100bp paired end reads as per manufacturer’s instructions.

### Whole Exome Sequencing Aligment and Variant Calling pipeline

FastQ sequencing files were aligned to the Hg19 reference genome using BWA-mem (version 0.7.7). Aligned sequencing files were converted to BAM files followed by sorting and indexing of reads using Samtools. Picardtools was used to mark duplicates and GATK (version 2.8) was used for local indel realignment. PicardTools, GATK (version 2.8) and FastQC were used to produce quality control metrics. SAMtools mpileup (version 0.1.19) was used to locate non-reference positions in tumour and germline samples. Bases with a Phred score of less than 20 or reads with a mapping quality <20 were omitted. MuTect (version 1.1.4) was used to detect SNVs, and results were filtered according to the filter parameter PASS. An SNV was considered a true positive if the variant allele frequency (VAF) was ≥ 5% and the number of reads in the tumour and germline at that position was ≥20. For insertion/deletions (INDELs), only calls classed as high confidence by VarScan2 and Scalpel were kept to avoid the risk of caller specific artefacts often observed with indel calling. Variant annotation was performed using Annovar (version 2016Feb01).

### Purity, ploidy and Copy Number (CN) estimation

The Sequenza package was used to derive copy number estimates for each sample. We obtained tumour purity and ploidy estimates using the probabilistic parameter search as suggested in the package manual. We included a quality control step in line with a recent publication (2) which examines the somatic SNV allele frequency distribution present in proposed regions of copy change. We found that all inferred copy states matched the expected allele frequency shifts suggesting our ploidy estimates were correct (allele frequency shifts includes peaks at 0.33 and 0.67 in trisomy regions, and 0.5 and 1 in regions of copy neutral LOH, following correction by tumour content).

### Calculation of Cancer Cell Fraction (CCF)

For each SNV, the CCF was calculated using a previously described formula based on the variant allele frequency (VAF), tumour purity and allele specific copy number (18). SNVs across all samples were pooled into 4 groups according to the MSH3/MSH6 mutation status of samples. The distribution of cancer cell fraction for SNVs and predicted neoantigens in each group was plotted as a density plot in R.

### Extraction of read length distribution of microsatellites

The MSIsensor package (version 0.6) was run on all samples using tumour and matching normal bam files as input. Default settings were adjusted to include a minimum homopolymer length of 6 for distribution analysis. Next the SciRoKo package (version 3.4) was used to identify all length 6-11 homopolymers within the exome using a bed file of genomic coordinates for the human Hg19 exome. MSIsensor distribution files were next filtered for exonic length 6-11 homopolymers and the resulting data used for downstream analysis.

### Identification of *MSH6*^F1088^ and *MSH3*^K383^ frameshift mutations

In order to accurately call frameshifts within *MSH6* and *MSH3* homopolymers we used the consensus of calls made using MSIsensor (version 0.6) and the variant calling pipeline described above. The MSIsensor derived microsatellite read length distribution of *MSH6* and *MSH3* homopolymers were extracted. The percentage of reads at each length was calculated for both microsatellites and tabulated (see Figure S11). To call a mutation a minimum of 5% of reads was required to show instability and with a minimum of 50 reads present. Next cases identified as mutated using this MSIsensor technique were checked against calls made using the Varscan/Scalpel pipeline described above. Any discrepancies were manually checked using IGV (Integrated Genomics Viewer v2.3) software.

### Shannon microsatellite diversity

The Shannon diversity index was calculated for all exonic length 6 to 11 microsatellites in each sample using the formula:

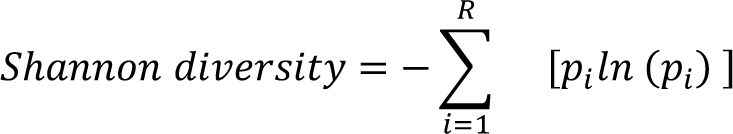

Where p_i_ = the proportion of total reads represented by the ith microsatellite length R= total number of read lengths present at a microsatellite

### Phylogenetic reconstruction

For tumours with more than 3 samples sequenced, the maximum parsimony method was used to infer phylogenies from the SNV calls. We used the Paup package (http://phylosolutions.com/paup-test/) and parameters as described previously(3). Briefly, SNV calls were converted into a binary matrix where 0 equals absence and 1 equals presence of a mutation, and rows relate to a biopsy or the normal sample, and columns each variant. The following methods were used for phylogenetic reconstruction: 1) the root function was used to root each phylogeny to the normal sample, 2) the hsearch function was used to perform a heuristic search of available trees and 1,000 of the shortest trees were output and examined, 3) the bootstrap function was used to randomly resample the data 10,000 times with replacement, with the proportion of each branch instance reported. The most parsimonious tree was reported for each case and in this data there was only ever one best solution. The .tre files generated were viewed using the FigTree software (http://tree.bio.ed.ac.uk/software/figtree/) and converted to PDF files.

For tumours with only 2 or 3 samples sequenced parsimony trees cannot be produced. For these cases the binary matrices were used to make simple inferences about clonality through shared mutation instances. Variants present in all samples were allocated as trunk mutations. For tumours with 3 biopsies, biopsy pairs with the most shared mutations were placed together on the same clade. Variants unique to each sample formed terminal branches (leaves).

### HLA typing and mutation calling

The Polysolver package (version 4) was used to perform haplotyping and mutation calling for HLA-A, HLA-B and HLA-C alleles. Germline and tumour sequencing data was supplied in the form of BAM files.

### Mutations in antigen processing machinery (APM) genes

Antigen processing machinery genes previously reported as undergoing mutation in MMRd cancers were identified from the literature (18). This created a gene list consisting of NLRC5, RFX5, TAP1, TAP2, CIITA and JAK1. Coding mutations in these genes (frameshift, non-synonymous SNVs or nonsense mutations) were retrieved from the annotated variant call files. Synonymous mutations were excluded.

### Neoantigen calling

Neoantigens were predicted using an established pipeline (Neopredpipe) utilising patient specific HLA haplotypes and the NetMHCpan prediction tool (4).

### Linear mixed effect model

To account for non-independence of multiple sampling per patient, a linear mixed effect model was created assessing the relationship between *MSH6/MSH3* frameshift status on total mutation burden. Individual variation in mutation burden between tumours was defined as a random effect and the presence of mutation in *MSH6* and/or *MSH3* microsatellites, age at diagnosis and tumour purity were defined as fixed effects. P-values were obtained by likelihood ratio tests of the full model with the effect of *MSH6/MSH3* status against the null model without the effect MSH6/MSH3 status. The model was created using the R package LME4 as follows: lmer(MT_burden ∼ MSH6_MSH3_status + age + tumour_purity+(1|Tumour_ID). Full results of the linear mixed effects model are provided in Supplementary table 4.

### Mutation signature analysis

Analysis of mutation signatures was performed using the package Sigprofiler (version 3.1). SNV and indel data were merged according to the *MSH6/MSH3* mutation status of samples resulting in the following 3 groups; samples wild-type for both *MSH6* and *MSH3*, samples with either the *MSH6^F1088fs^* or *MSH3^K383fs^* and samples with both *MSH6^F1088fs^* and *MSH3^K383fs^*. Three de-novo SBS signatures were extracted and their 96-channel trinucleotide context was plotted. The percentage contribution of each signature according to *MSH6/3* meta-groups was further plotted. Similar analysis was performed for indel and double base mutations.

### Immune dN/dS

Immune dN/dS, defined as the portion of the genome exposed to immune recognition, was calculated using SOPRANO(5) (the code is available at github.com/luisgls/SOPRANO). It estimates dN/dS values in a target region (ON-target) and in the rest of the proteome (OFF-target) using a trinucleotide context correction (SSB192). Here, we have used genomic regions that translate to peptides that bind HLA-A0201 allele as the target region (ON). Only genes with a median expression of more than 1FPKM were used according to the human expression atlas data (downloaded on Oct 18/2018). The file used as target region can be obtained from github.com/luisgls/SOPRANO.

### MOBSTER clonal deconvolution and mutation rate analysis

In order to retrieve in-vivo mutation rates estimates we developed a simple pipeline around MOBSTER, a recently developed computational method that can perform tumour subclonal deconvolution by integrating population genetics and Machine Learning(6). This method is able to retrieve an estimate of the tumour mutation rate (μ) from the tail of neutral mutations. To run it, we first pooled somatic variants and absolute copy number alterations (CNAs) generated as detailed above. We then used a computational method to map somatic SNVs on top of copy number segments, and assess the consistency between tumour purity, ploidy and CNAs. We restricted our analysis to SNVs and dropped indels because the Variant Allele Frequencies (VAFs) estimates for SNVs are more reliable for assessing the quality of the calls and performing deconvolutions. All the samples we analysed passed our quality check process.

We then assessed the overall percentage of CNA segments that span the tumour genome, and considered the copy state of each segment. This confirmed that the largest chunk of tumour genome is in heterozygous diploid state, with a single copy of the major and minor alleles, which is expected from colorectal cancers with microsatellite instability(7). For this reason, we retained only SNVs mapping to diploid segments, which harbour less noise compared to mutations that map to more complex tumour karyotypes. With pooled diploid SNVs we proceeded to run tumour subclonal deconvolution, using raw VAFs and MOBSTER. The tool was run to search for tumours with up to 2 subclones, with an optional neutral tail; model selection for the number of clonal populations (k>0) and the tail was performed using the routines available in MOBSTER. MOBSTER could estimate the different mixture of cancer subpopulations in each one of the bulk samples, as well as the neutral tail of somatic SNVs that accrue inside each of the subclonal expansions, if present. Across data for cancer UCL_1014 we observed monoclonal populations (k=1) with a neutral tail, therefore concluding that these data lack evidence of ongoing subclonal positive selection, consistently with patterns of colorectal cancer evolution observed earlier (8).

Parameters of the fit for the Power Law Type-I tail available in MOBSTER were then used to retrieve the tumour mutation rate >0. This quantity is canonically expressed in time units of tumour cell doublings - i.e., considering discrete time-evolution steps in which all tumour cells divide synchronously - and depends on the size of the analysed tumour genome. To make it comparable across multiple samples of the same patient, and to account for the fact that we used whole-exome data, we normalised by the size of the diploid exome regions in each biopsy of every patient. These gave us the point estimates reported in the Main Text.

We then sought out to build a Confidence Interval (CI) for μ, adopting a non-parametric bootstrap procedure(9). In practice, we bootstrapped with repetitions from the mutations available in each sample and built n=200 datasets per patient; then we re-run the MOBSTER analysis conditioning on retrieving the putative monoclonal architecture (k=1) identified in the main run, and re-computed the normalised values for the bootstrap estimate of μ. With the distribution of bootstrapped μ values, we built a percentile CI corresponding to an α-level of 5% by taking the 2.5% and 97.5% empirical quantiles.

### Multiplex Immunofluorescence

A panel consisting of MSH6, CD20, FOXP3, CD4, PANCK and CD8 were selected for the multiplex immunofluorescence assay. Primary antibody details are provided in supplementary table S5. The Opal (Akoya) multiplex immunofluorescence automation kit was used which includes HRP conjugated secondary antibody, Opal fluorophores, DAPI stain, antibody diluents and blocking buffers. The manufacturer’s protocol was followed and immuno-staining performed using the Leica Bond RX autostainer (Leica Biosystems).

### Monoplex optimization

To optimize labelling of each marker, monoplex slides were created where tissue sections were labelled with each primary antibody on separate slides. Each primary antibody was assigned an opal fluorophore. Monoplex slides were processed with appropriate number of antibody stripping steps before and after staining reflecting the eventual multiplex sequence. Monoplex slides were imaged using the Vectra 3.0 fluorescence microscope and signal counts assessed using the Inform software.

### Autofluorescence slide

A representative tumour section was labelled with Pan-CK primary antibody and without opal fluorophore to assess levels of background autofluorescence.

### Library development

To create a spectral unmixing library, slides were stained with the most abundant marker (Pan-CK) and each opal fluorophore individually, resulting in 6 library slides. Library and autofluorescence slides were imaged on the Vectra 3.0 using all 5 epi-fluorescence filters (DAPI, FITC, Cy3, Texas red, Cy5). The spectral unmixing library was developed using the Inform software.

### Multiplex assay development

The multiplex assay was run using the optimized conditions developed during monoplex optimisation. Timings, temperature settings and reagent concentrations for each step are detailed in Supplementary table 5. The following steps were performed on the Leica BondRx autostainer:

1. Deparaffinization using Bond dewax solution
2. Antigen retrieval solution using Bond ER1 or ER2 solution
3. Blocking buffer
4. Primary antibody incubation
5. Opal polymer HRP incubation
6. Opal fluorophore incubation
7. Stripping of antibody complexes using Bond ER1 or ER2 solution
8. Repeat steps 2-7 until all primary antibodies applied
9. DAPI counterstain

Following immunostaining steps above the slides were cover-slipped manually using Diamond antifade mountant (Invitrogen) and imaged using the Vectra 3.0.

### ORION cell segmentation workflow

In this study we developed ORION (FluORescence cell segmentatION), a cell segmentation workflow for multispectral immunofluorescence imaging (see figure S5 for overview). ORION uses an ellipsoidal model(10) to identify individual cells, exclude noise and non-cell objects. To this end, initially, the unmixed spectral signatures undergo a Gaussian filter with a 5 × 5 kernel in order to remove small artifacts. Subsequently, an adaptive thresholding method is applied for the selection of the optimal threshold value for each pixel within its local neighbourhood(11). This requires mean filtering and estimation of the local threshold based on the mean neighbourhood pixel intensity. In the resulting *M* binary image, morphological operations consisting of erosion, dilation and removal of small elements are applied, in order to suppress small artifacts.

For the separation of touching cells an improved ellipsoidal modelling approach is performed. Initially, we estimate the distance transformation of the binary image *M* of *p* pixels that represents the connected cells and we estimate the regional maxima of this. Given that the number and location of local maxima correspond to those of nuclei, we reject the touching maxima. The remaining maxima comprise the list of candidate seeds. Then, based on the hypothesis that cells can be spatially modelled as ellipsoids *E*_c_, the pixels of cells are then modeled using a Gaussian distribution. More specifically, a Gaussian mixture model is applied with the number of clusters c being equal to that of candidate seeds and the mixture parameters, namely the mean and variance, are estimated using the expectation-maximization (EM) algorithm. For the initialization of the EM algorithm the *k*-Nearest Neighbor classification using Euclidean distance as the distance metric is used in order to estimate the initial parameters. The EM is an iterative method consisting of two steps: (i) expectation, which computes the likelihood with respect to the current estimates and (ii) maximization (Eq. 1), which maximizes the expected log likelihood (Eq. 2) as follows:

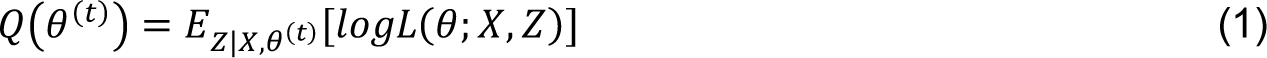

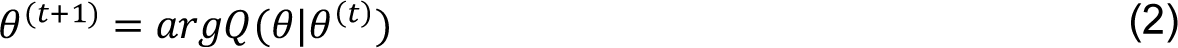

where *Q* is the expected values of the log likelihood function θ, *X* is the pixel coordinates, *Z* is the latent variables and θ^(t)^ is the current parameters.

Having estimated the ellipsoidal model of cells, we need to reject any erroneous seeds from the candidate list and re-estimate the ellipsoidal models for the remaining seeds. For this study, we developed a new fitness validation criterion taking into account the overall combination of ellipses of candidate seeds. More specifically, the proposed criterion aims to keep the cells well-separated and takes into account the binary areas that are included in the estimated ellipses, the total area of the extracted ellipses, as well as the background area that is included in the estimated ellipses and the overlapping parts of the ellipses of the touching cells. Subsequently, we introduce an intensity-based parameter *W*_I_ based on the intensity variance of each estimated ellipse aiming to separate the touching cells with different intensity. In the case that the estimated ellipses fit perfectly to the binary mask *M* the value of fitness function tends to be equal to 1. To this end, the number of candidate seeds and the estimated ellipsoidal components are defined by maximizing the following fitness degree of the 2D cell data mask:

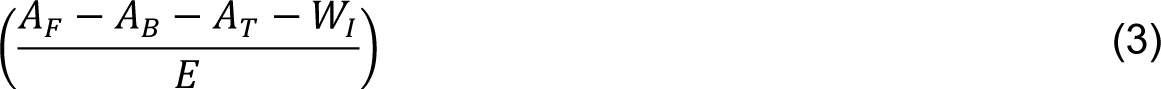

where the total area covered by the estimated ellipses is *E* = ∑_*p∈M*_ *E*_c_(*p*), the foreground area of the binary image *M* that is included in the estimated ellipses is A_F_ = ∑_*p*=1_ *M*(*p*)*E*(*p*), the area of the background area of the binary image *M* that is included in the estimated ellipses is A_B_ = ∑_*p*=1_ [1 − *M*(*p*)]*E*(*p*) and the overlapping parts of the ellipses of the touching cells for the total number of the identified ellipses is defined as 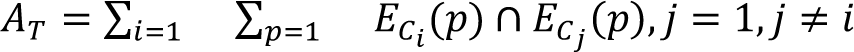. The final segmentation of the clustered cells is performed by applying Bayesian classification which assigns each pixel *p* to cluster c_i_ with the maximum posterior probability.

To validate the performance of ORION, we conducted tests using two datasets and compared with 7 independent state-of-the-art cell segmentation approaches. The two datasets used for this evaluation were a publicly available dataset (dataset A) and a subset of the multispectral IF images of MSI tumours created in this study (Dataset B). Dataset A included 48 fluorescence images of 1831 cells with nuclei labelled using the Hoechst stain(12). Dataset B consisted of 400 cells from multiplex IF images selected randomly from our study dataset. To validate the efficiency of ORION we used Jaccard Similarity Coefficient (JSC), as well as Dice false positive (Dice FP) and Dice false negative (Dice FN) values in order to measure over-segmentation and the under-segmentation respectively. Furthermore, Hausdorff distance and mean absolute contour distance (MAD) were used to evaluate the contour of detected cells. Moreover, we estimated the True Detected Rate in order to determine the ratio of segmented cell numbers to the total number of annotated cells present. Using ORION, cell segmentation of different markers was carried out based on DAPI nuclear staining or combining DAPI and cytoplasmic staining due to lack of clear cell boundaries (e.g. CD4+ cells). Since both tumour cells and immune cells may express MSH6, we used co-localisation of MSH6 expression with the epithelial marker pan-CK to identify tumour cells and immune markers (CD8, CD4 and CD20) to identify specific immune cell populations. The experimental results (Supplementary table 6) show that ORION outperformed seven state-of-the-art approaches that have been used in the past for cell segmentation. Finally, the True Detected Rate for the two datasets was estimated to be equal to 98.9%.

Following validation of ORION, we next ran the workflow on the full dataset of 194 multispectral image tiles from 27 tumours. We performed neighbourhood analysis(13) to quantify the number of immune cells of each subtype within the vicinity of MSH6 proficient and deficient tumour cells. Image tiles were classified as MSH6 proficient or deficient based on the expression status tumour cells. We then used the localized segmented cell centres of tumour cells and a radius *R = 100um* in order to identify the spatial associations with immune cells. We chose this radius as a biologically relevant distance for interaction between tumour and immune cells. We counted the number of each type of immune cell identified within this radius. For each tile we reported the sum total of immune cells of each subtype identified from neighbourhood analysis.

### Genomics England 100,000 Genomes CRC Dataset

#### Whole Genome Sequencing, variant calling, purity and ploidy estimation

Whole genome sequencing data were generated through a standardised, Clinical Pathology Accredited, workflow as part of the Genomics England 100,000 genomes project(14). Briefly, the sequencing data was aligned to human genome GrCh38 using the Illumina iSAAC aligner and was then subjected to extensive variant calling and quality control processes. The basis of this is the Strelka variant caller plus artefact filtering using a project-wide panel of normal, and population-based filtering using the aggregated gnomAD dataset. We also scrutinised the resultant SNV calls taking a minimum depth cutoff of 50X. We used the package Sequenza to derive tumour purity, ploidy and copy number estimates for each sample as described above. In total 992 colorectal cancers were identified for study using the V8 data release (available as of 11/2019).

#### Identification of MSI cancers

Cancers with microsatellite instability were detected using the MSIsensor (version 0.6) package and validated using two methods. Firstly, using the mutational signature analysis available as part of the V8 release, we confirmed the enrichment of a mismatch repair deficient SBS mutation signature. As further validation, cases with available pathological data were confirmed to be mismatch repair deficient by immunohistochemistry (histology validation set, n = 101, 98% classification accuracy). Tumours with discernible pathogenic POLE or POLD1 exonuclease domain mutations were excluded. The resulting cohort of 217 cases were used for downstream analysis and referred to here as the GEL CRC MSI cohort.

#### Identification of primary MMR defect

Germline and somatic mutations in the MMR genes (MLH1, PMS2, MSH2, MSH6 and MSH3) were identified by searching the relevant Genomics England main programme tiering data for tier 1 pathogenic mutations. To identify tumours with MLH1 promoter methylation, the presence of somatic BRAFV600E was used as an indicator. This approach is in keeping with current clinical guidelines where it is recognized that amongst MMRd colorectal tumours the presence of somatic BRAFV600E mutation associates strongly with MLH1 promoter methylation (7).

### Mutation frequency of *MSH6/MSH3* microsatellites compared with other length 8 coding microsatellites

Since *MSH6* and *MSH3* contain a C8 and A8 coding homopolymer respectively, we were interested to compare the frequency with which these sites are mutated across the cohort compared to other length 8 homopolymers of the same nucleotide base. The genomic coordinates of all length 8 exonic microsatellites were obtained using the SciRoKo package. The mutation status of all length 8 coding microsatellites and the base affected were extracted from the variant call files by filtering for the genomic coordinates of length 8 coding microsatellites. The percentage of cases with frameshift mutation in C:G or A:T microsatellites were calculated separately and compared to the mutation frequency observed for *MSH6* (C8) and *MSH3* (A8) microsatellites respectively. Comparisons were made using the Chi squared test.

### Multiple linear regression model

To determine the contribution of *MSH6^F1088fs^* and *MSH3^K383fs^* frameshifts to mutation burden in MSI CRCs, multiple linear regression modelling was performed. The presence or absence of frameshift mutation in the coding homopolymers of *MSH6* and *MSH3* together with mutation status of coding homopolymers in a further 21 genes were used as independent variables in the model. The list of 21 genes were those reported as recurrently mutated in MSI CRC(15) and consisted of *RFX5, MBD4, AIM2, ACVR2A, DOCK3, TGFBR2, GLYR1, OR51E, CLOCK, CASP, JAK1, TAF1B, BAX, MYH11, HPS1, SLAMF1, HNF1A, RGS12, ELAVL3, SMAP1, SLC22A9*. We also included tumour purity and age at diagnosis in the model to account for potential confounding. There was no difference in estimated tumour purity between MSH6 and MSH3 mutated groups (one-way anova p=0.742). The model was created using the lm function in R and results were plotted as a volcano plot with regression coefficients of contribution to mutation burden versus -log_10_ p value of the t-statistic. We also ran the model using just *MSH6* and *MSH3* frameshifts as independent variables and obtained estimates of the contribution of these frameshifts individually and combined on total mutation burden. Results of the model output are provided in supplementary table S1.

### Identification of *MSH6/MSH3* coding mutations outside of length 8 homopolymers

Variant call files were searched for all coding *MSH6* and *MSH3* mutations. Data were extracted and tabulated according to the frequency and type of mutation (figure 1B and supplementary table S2).

### Mutation burden analysis

Total mutation, SNV and indel counts for each sample was measured using data from variant call files. Both synonymous and non-synonymous SNVs were included. Violin and waterfall plots were generated in R using the ggplot package.

### Analysis of MSI cases with confirmed primary MLH1/PMS2 (MutL) deficiency

In order to confirm that differences in the primary cause of MMR loss in samples was not confounding results, we restricted our analysis to cases with confirmed MLH1/PMS2 (MutLα) deficiency. We identified tumours with somatic *BRAF*^V600E^ mutation (indicating *MLH1* promoter methylation) and also samples with tier 1 pathogenic germline *MLH1* or *PMS2* mutations. Somatic BRAF^V600E^ mutations were identified from variant call files and germline mutations from the Genomics England main programme tiering data. Violin plots for total mutation, SNV and indel burden were plotted on this subset of the cohort as previously described.

### TCGA dataset

MSI cancers within the TCGA dataset were identified from a previous study(15). Variant calls for these tumours were downloaded from the National Cancer Institute Genomics Data Commons portal (https://portal.gdc.cancer.gov/repository). Cases with frameshift mutation in the *MSH6* and *MSH3* coding microsatellites were identified from supplementary data provided in(15). Tumours with discernible mutations in *POLE* and *POLD* exonuclease domains were excluded(16). This resulted in a MSI cohort consisting of the following tumour types, colorectal (n=48), uterine (n=67), stomach (n=63) and esophageal (n=3). Mutation burden plots were created in R using the ggplot package according to the *MSH6/MSH3* mutation status of tumours. To analyse neoantigen counts data were obtained from Lakatos et al (17). Briefly tumour purity and ploidy were estimated using ASCAT on Affymetrix SNP array data. Samples with purity below 0.4 and ploidy above 3.6 were excluded. This reduced the cohort size to 117 samples (colorectal = 34, uterine=56, stomach=27). Neoantigens were predicted using the Neopredpipe pipeline as detailed in(^4^). To analyse MSH3 and MSH6 RNA expression levels, raw RNA counts were obtained and transformed to FPKM (fragments per kilobase million) and then converted to TPM (transcripts per kilobase million) values using the formula tpm = fpkm/sum(fpkm) * 10^6^. RNA expression data was available for 127 samples (colorectal n=42, uterine n=56, stomach n=29).

### Mathematical model of the effect of stochastic mutation rate switching on tumour growth

Our model was based on our previous stochastic branching process modelling of tumour growth and neoantigen accumulation(17). The model simulated tumour growth where each cell can either (i) die with a probability inversely proportional to their fitness or (ii) divide into two daughter cells that accumulate novel mutations according to their respective mutation rate. Cells in hyper-mutated and ultra-hyper-mutated state gain a number of mutations in each division sampled from a Poisson distribution with parameter (mutation rate, μ) 6 and 120, respectively. New mutations are either (i) neutral with no effect on cell fitness; (ii) antigenic, decreasing the cell’s fitness; (iii) immune escape mutations that eliminate immune predation and therefore nullify antigen-induced fitness decrease; (iv) lethal, irreversibly decreasing cell fitness regardless of immune escape (Fig. 4A). The probability of a given mutation being non-neutral is defined by p(antigen), p(escape), p(lethal), respectively. Note that these mutation types are non-exclusive and a mutation can be for example both antigenic and lethal (though with only a small probability). In addition, at each division the daughter cells may undergo mutation rate switching with probability β. β=0 corresponds to no switching (mutation rates remain constant) while β>1/100 represent frequent switches to or from ultra-hyper-mutated state. Each tumour was initiated with a homogeneous population of 100 tumour cells all in either hyper-mutated or ultra-hyper-mutated state; and simulated until elimination (no tumour cells left) or until reached detectable size (>100,000 cells).

### Microsatellite diversity

We encoded the mutation status of the microsatellite locus as an integer: 0 represented a wild-type allele, −1/+1 a single deletion/insertion, and so on. Upon division, daughter cells inherited the mutation status of their ancestor. Every new mutation had a probability, p(ms), to affect the microsatellite: if they did, the state of the locus was changed from N to N+1 or N-1 with equal probability. At the end of the simulation, the mutation status of all cells was read out and the total Shannon diversity of the population was computed in R (using the package entropy).

### Growth time

We defined the start of the “growth period” as the last time-point when the population count went below 20 (immune escaped) cells. The final time was the time-point when the population reached 100,000 cells. Growth time was computed as T(final) – T(growth-start). We chose this measure over T(final) as the latter had a very high uncertainty due to the variable time lineages spent before probabilistically acquiring immune escape and initiating unimpeded growth.

### Parameter values

The following default parameter values were used in all simulations, unless indicated otherwise (e.g. a range of p(lethal) values in Fig. 4F): Neoantigen probability (p(antigen)): 0.1; immune escape probability (p(escape)): 10^-6^; lethal mutation probability (p(lethal)): 5*10^-4^; microsatellite-shifting rate (p(ms)): 10^-3^; immune-related selection coefficient (s): −0.8 (representing moderate selection).

## Statistical analysis

The Kruskal-Wallis test was used to test for a difference in distribution of 3 or more groups with post-hoc pair-wise comparisons performed using the Wilcoxon unpaired test. A P value of less than 0.05 was considered significant. Multiple linear regression was performed using the Lm package and linear mixed effect modelling performed using LMER package in R. For correlation analysis Pearson and Spearman rank correlation was used to assess for linear and monotonic relationships respectively. The Chi-squared test was used to compare the distribution of categorical variables. Statistical analyses were performed in R (version 3.6.2).

## Supplementary figure legends

**Figure S1.**
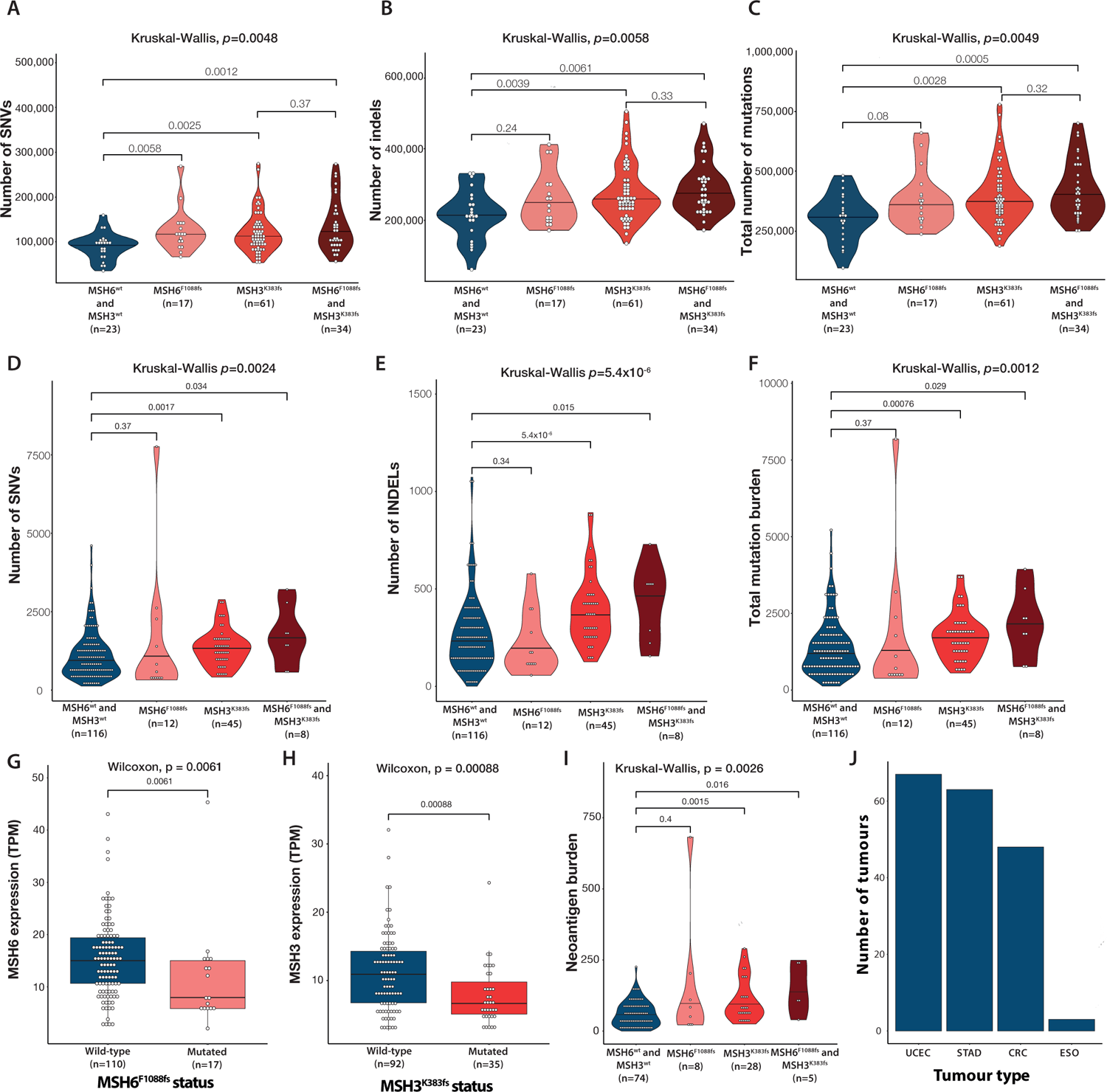
Mutation burden is increased in presence of secondary MSH6^F1088fs^ and MSH3^K383fs^. (A-C) GEL MSI CRC cohort subset for cases with confirmed MutL (MLH1/PMS2) loss as primary cause of MMRd. Violin plots show SNV, indel and total mutation burden in tumours according to presence of secondary MSH6 F1088 and MSH3 K383 frameshifts. (D-J) TCGA MSI validation cohort (D-F) Violin plots display SNV, indel and total mutation burden according to presence of secondary MSH6 F1088 and MSH3 K383 frameshifts. (G-H) RNA expression of MSH6 and MSH3 is reduced in tumours with MSH6 and MSH3 frameshifts respectively. (I) Neoantigen burden according to MSH6 F1088 and MSH3 K383 frameshifts. (J) Number of tumours according to tumour type making up the cohort. UCEC = uterine corpus endometrial carcinoma, STAD = stomach adenocarcinoma, CRC = colorectal adenocarcinoma and ESO = esophageal adenocarcinoma.

**Figure S2.**
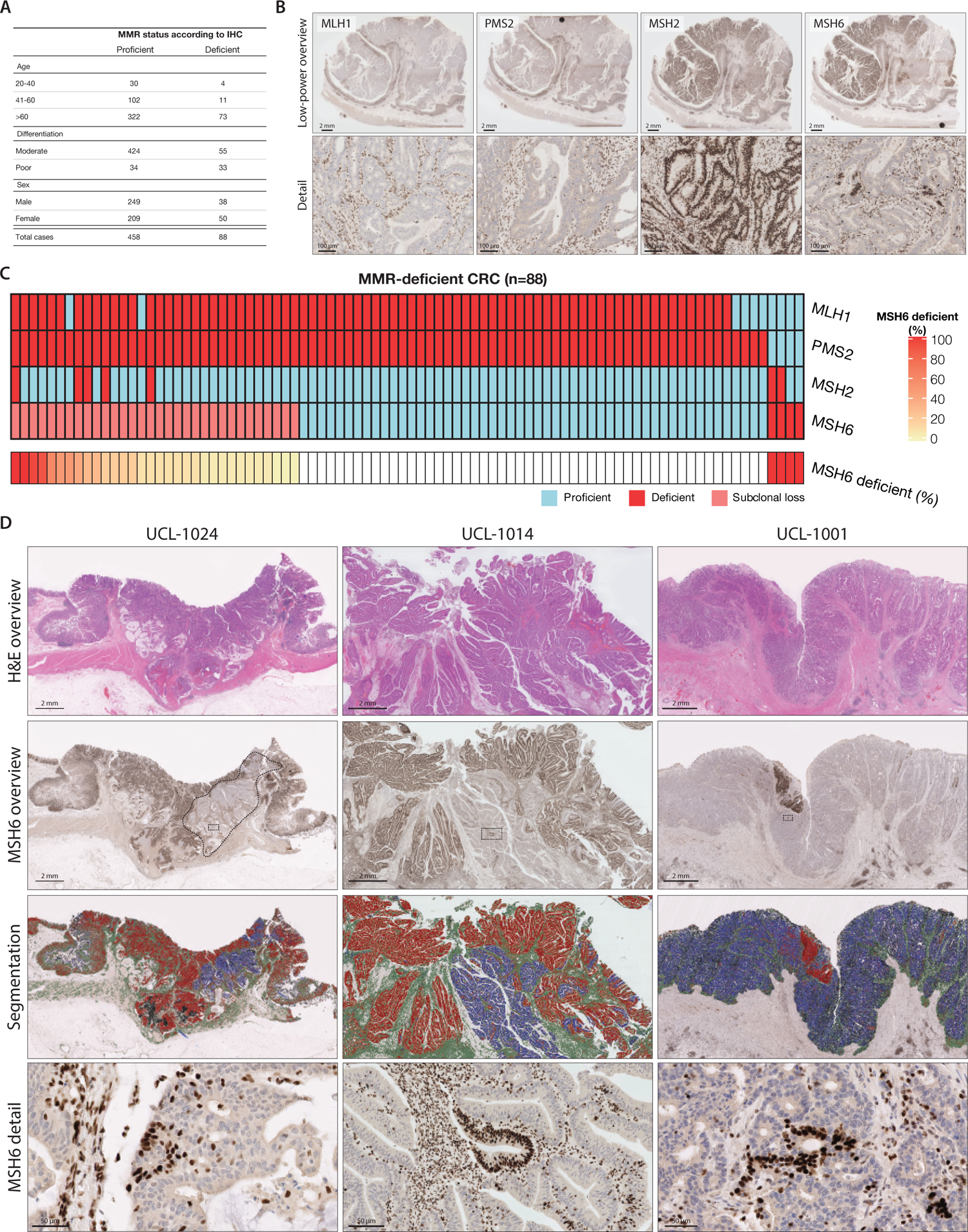
MSH6 immunolabelling reveals nested proficient reversion subclones within deficient regions. (A) Summary table for the cohort of 546 consecutive CRCs used in this study. (B) MMR IHC of polypoid tumour with complete loss of MLH1/PMS2 and subclonal loss of MSH6. High power detail image shows islands of MSH6 reversion (arrowheads) within MSH6-deficient subclone. (C) Breakdown of the pattern of MMR protein loss in 88 MMRd tumours. 32 tumours had subclonal MSH6 loss and the estimated percentage of deficient tumour cells observed is displayed in the heatmap. (D) Three example tumours with MSH6 subclonal loss. For each tumour H&E staining overview, MSH6 immunohistochemistry overview, IHC segmentation (red is MSH6-proficient, blue is MSH6 deficient), and detail images are shown.

**Figure S3.**
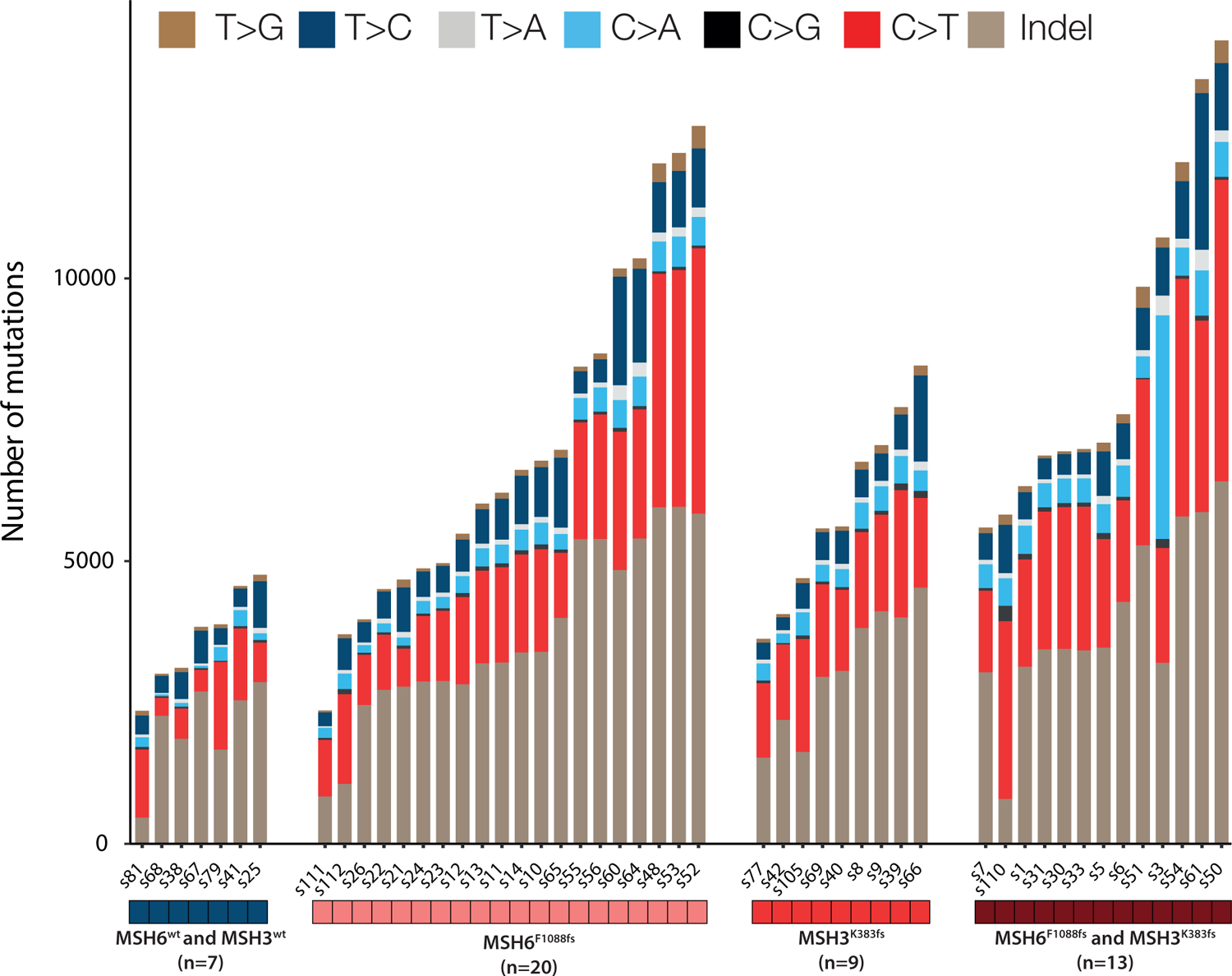
Mutation bias on a per-sample basis with absolute numbers of each mutation type.

**Figure S4.**
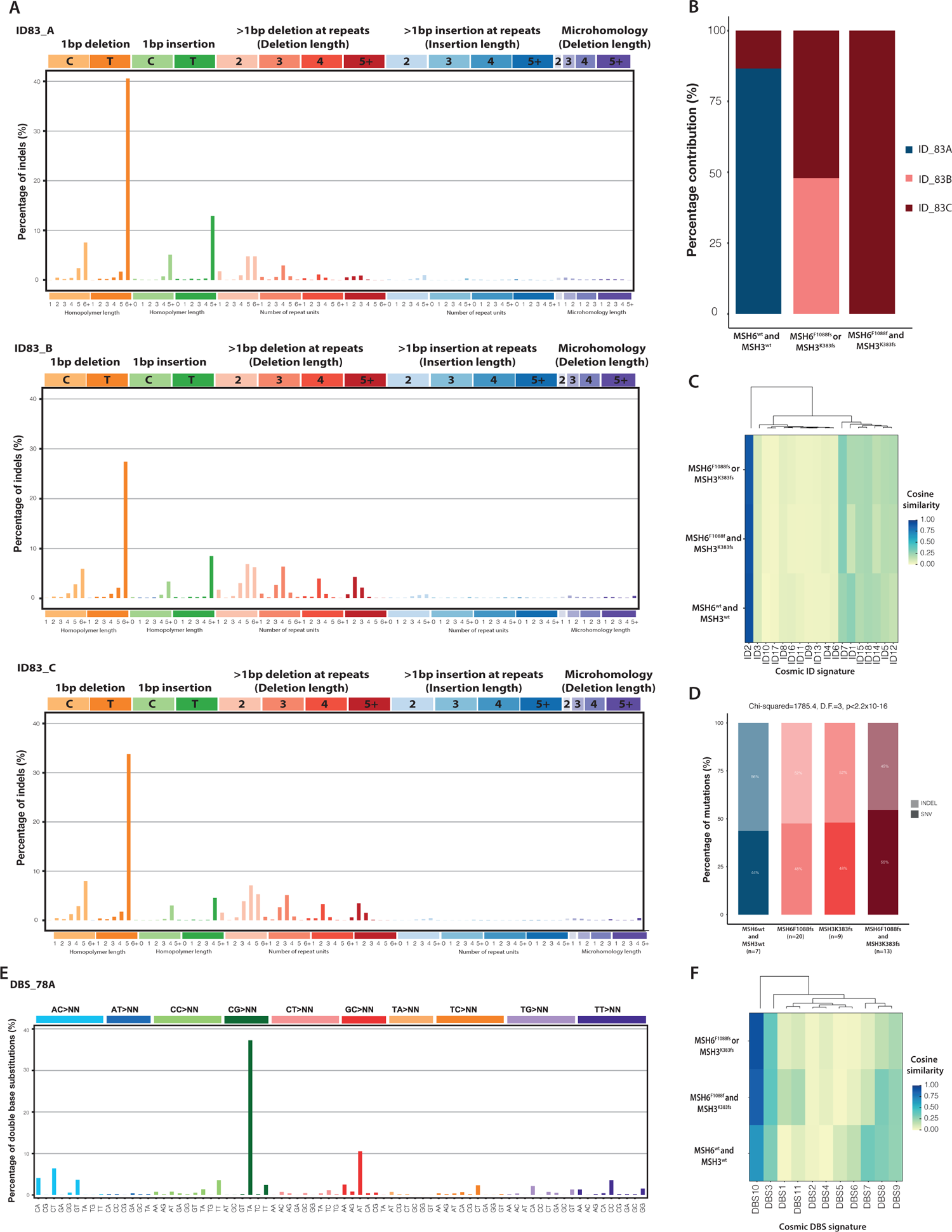
Indel and Double Base Signatures (DBS) (A) Extracted de-novo indel signatures A,B & C displayed according to contribution of mutations in 83 category format. (B) Relative contribution of each extracted de-novo indel signatures in samples grouped by absence or presence of MSH6^F1088^ and MSH3^K383^ frameshifts. (C) Cosine similarity of de-novo indel signatures to existing COSMIC indel signatures. (D) Proportion of substitutions and indels according MSH6^F1088^ and MSH3^K383^ frameshifts. (E) Extracted de-novo DBS signature displayed according to contribution of mutations in 78 category format. (F) Cosine similarity of de-novo DBS signature to existing COSMIC DBS signatures.

**Figure S5.**
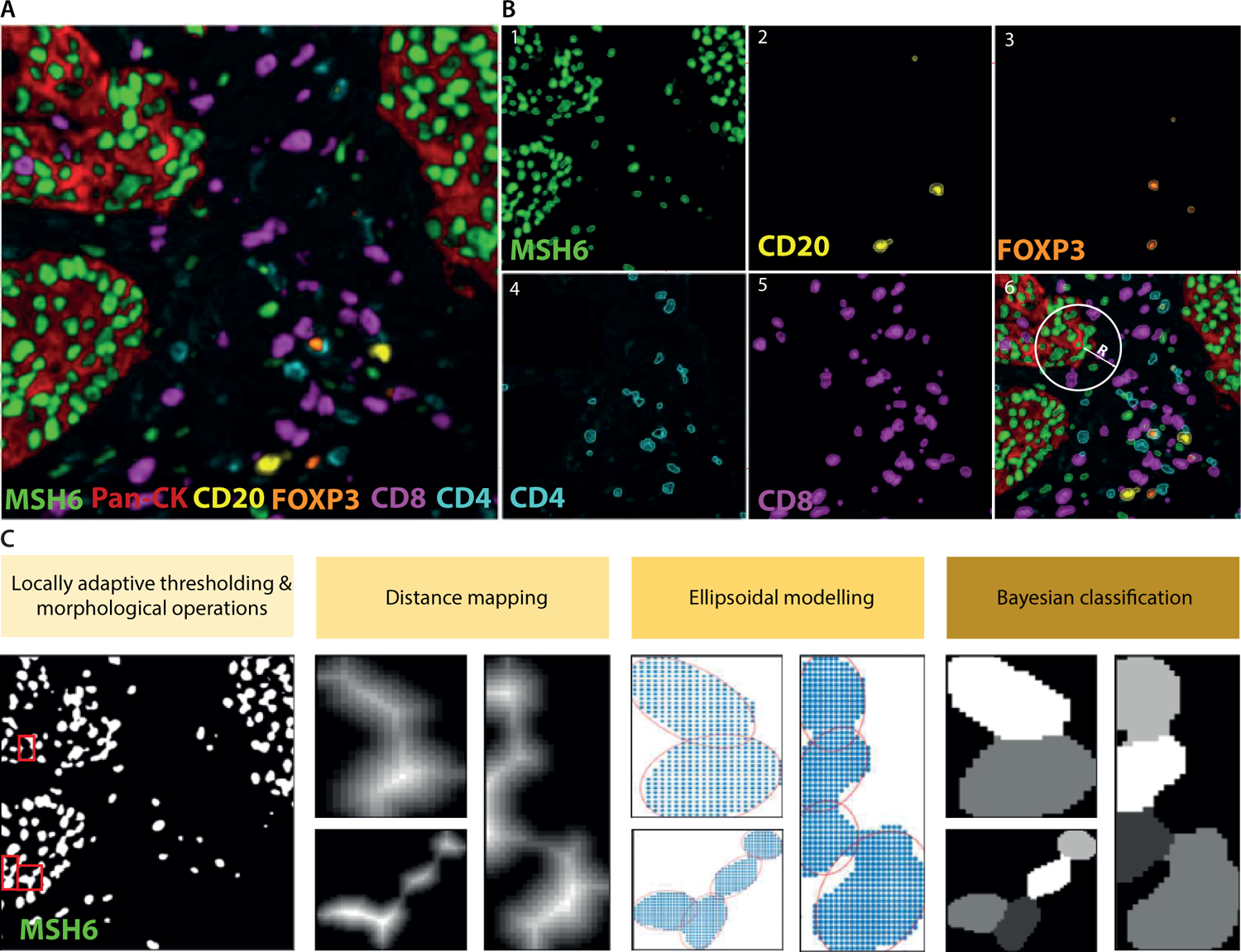
ORION (FluOResence cell segmentatION) workflow. | (A) Multiplex IF image of example tumour labelled for MSH6, PanCK, CD8, CD4, CD20 and FOXP3. (B) Isolation of MSH6, CD20, FoxP3, CD4 and CD8 spectral signals. (C) Example of the ORION main workflow steps. The workflow includes locally adaptive thresholding of isolated spectral signals, estimation of distance maps and local maxima, ellipsoidal modelling of cells and Bayesian classification for the identification of cells. Neighbourhood analysis is used for the identification of tumour-immune interactions estimating the number of immune cells within radius R of each MSH6 proficient or deficient tumour cell (shown in B6).

**Figure S6.**
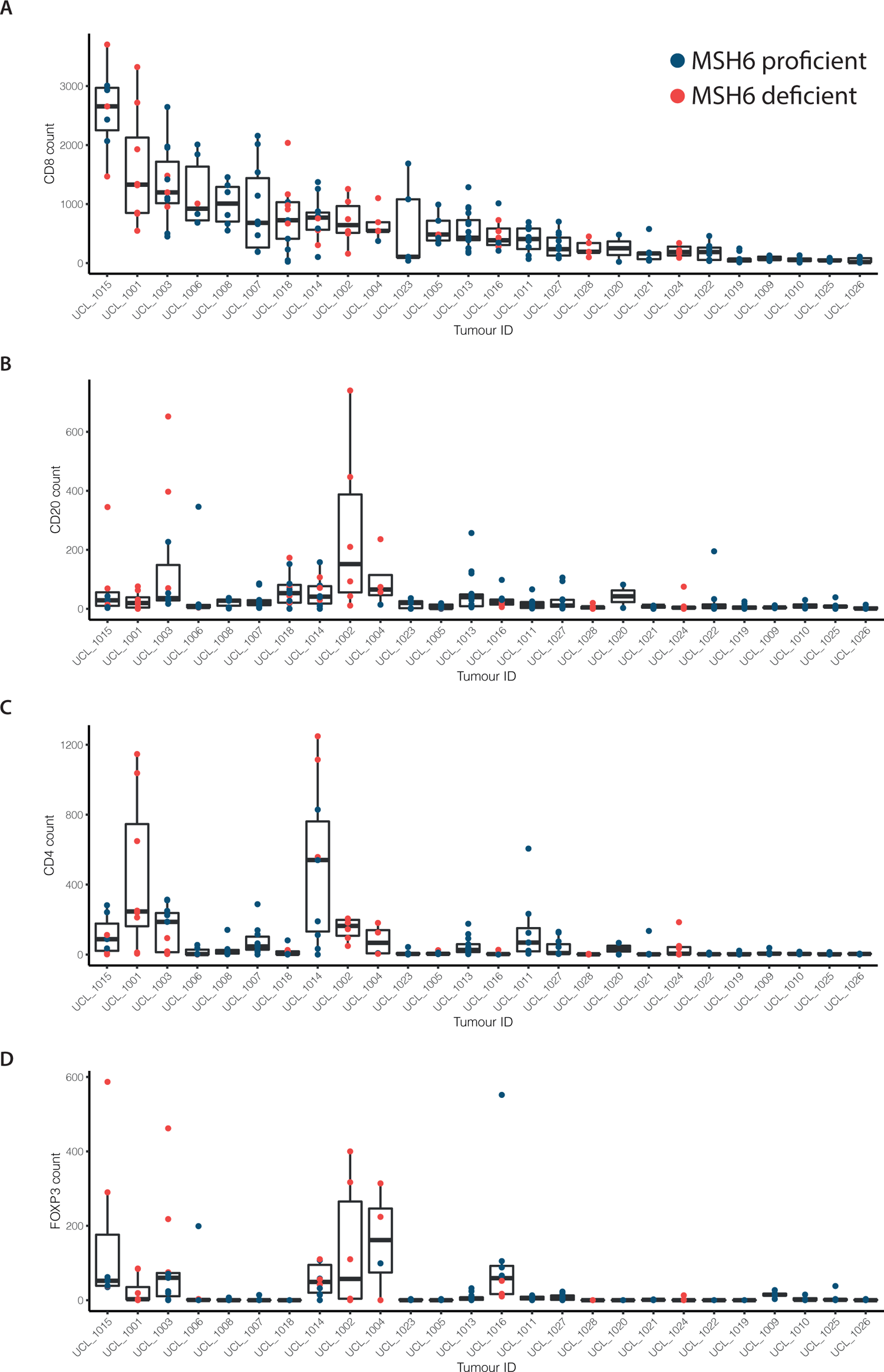
Immune infiltration in tumours according to MSH6 status of imaged tile. (A-D) Immune infiltration levels for CD8, CD20,CD4 and FOXP3 cells in tumours per imaged tile. Tiles are coloured according to the MSH6 expression status of tumour cells.

**Figure S7.**
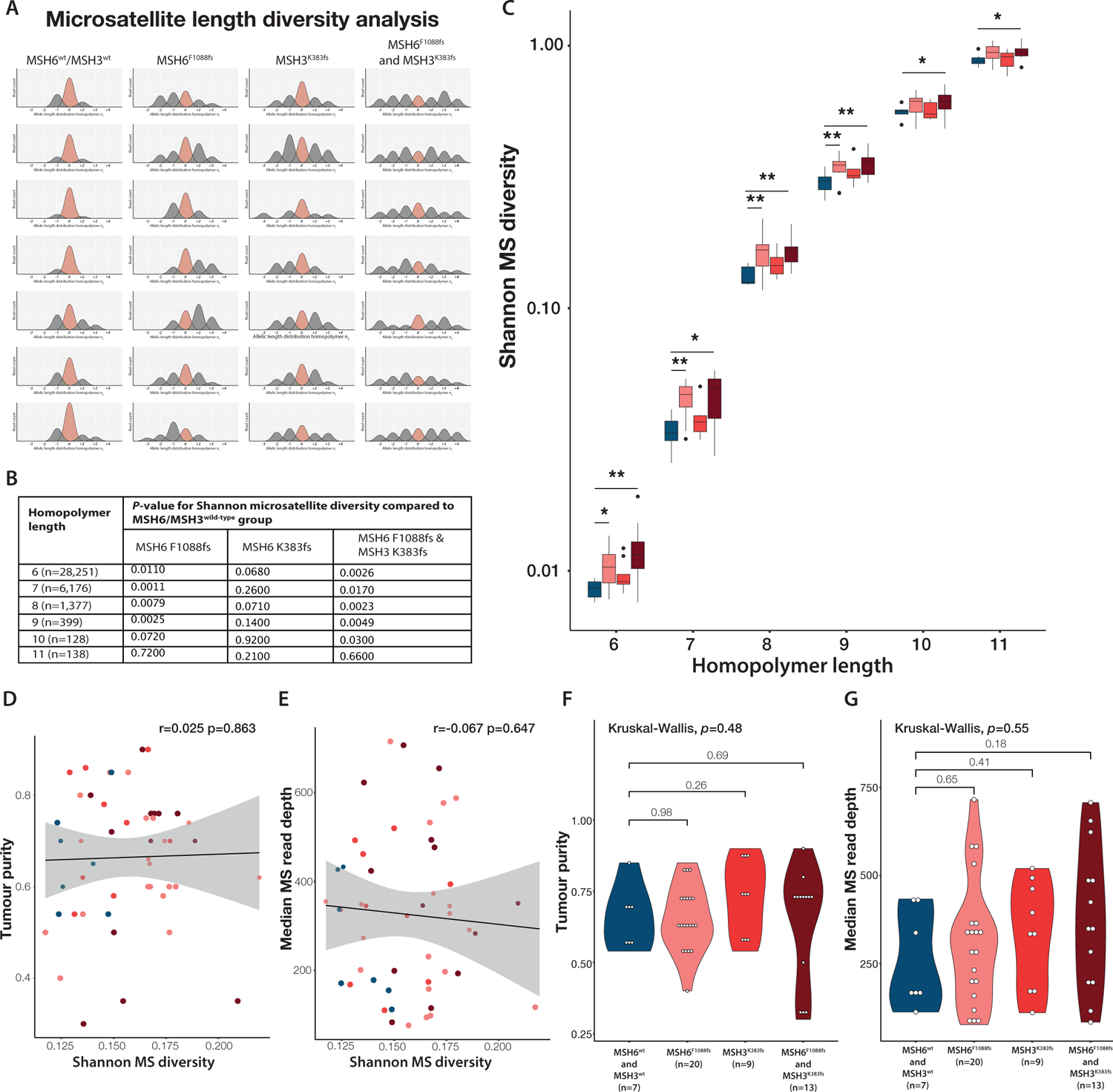
Microsatellite diversity in tumour samples according to homopolymer length. (A) Cartoon illustrating microsatellite length diversity analysis. We hypothesised that increasing clonal mutation burden would be reflected in increasing clonal diversity. In this analysis, microsatellites are used as lineage tags to interrogate population structure. MSH6 and MSH3 frameshift status from left to right and seven arbitrary homopolymers from top to bottom. Plots show read count for each microsatellite length, beige is the wild-type reference allele and grey are deviations from the reference. Cartoon shows progressive population heterogeneity with *MSH6* and *MSH3* frameshifts. (B-C) Homopolymer length versus Shannon microsatellite diversity in samples grouped according to MSH6 and MSH3 frameshift status. *MSH6* and *MSH3* frameshifts result in increased microsatellite diversity, but effect size decreases at longer homopolymer lengths. (D-E) No correlation between Shannon MS diversity and tumour purity or median microsatellite read depth. (F-G) No difference in tumour purity or read depth at microsatellite sites between samples according to MSH6/MSH3 grouping.

**Figure S8.**
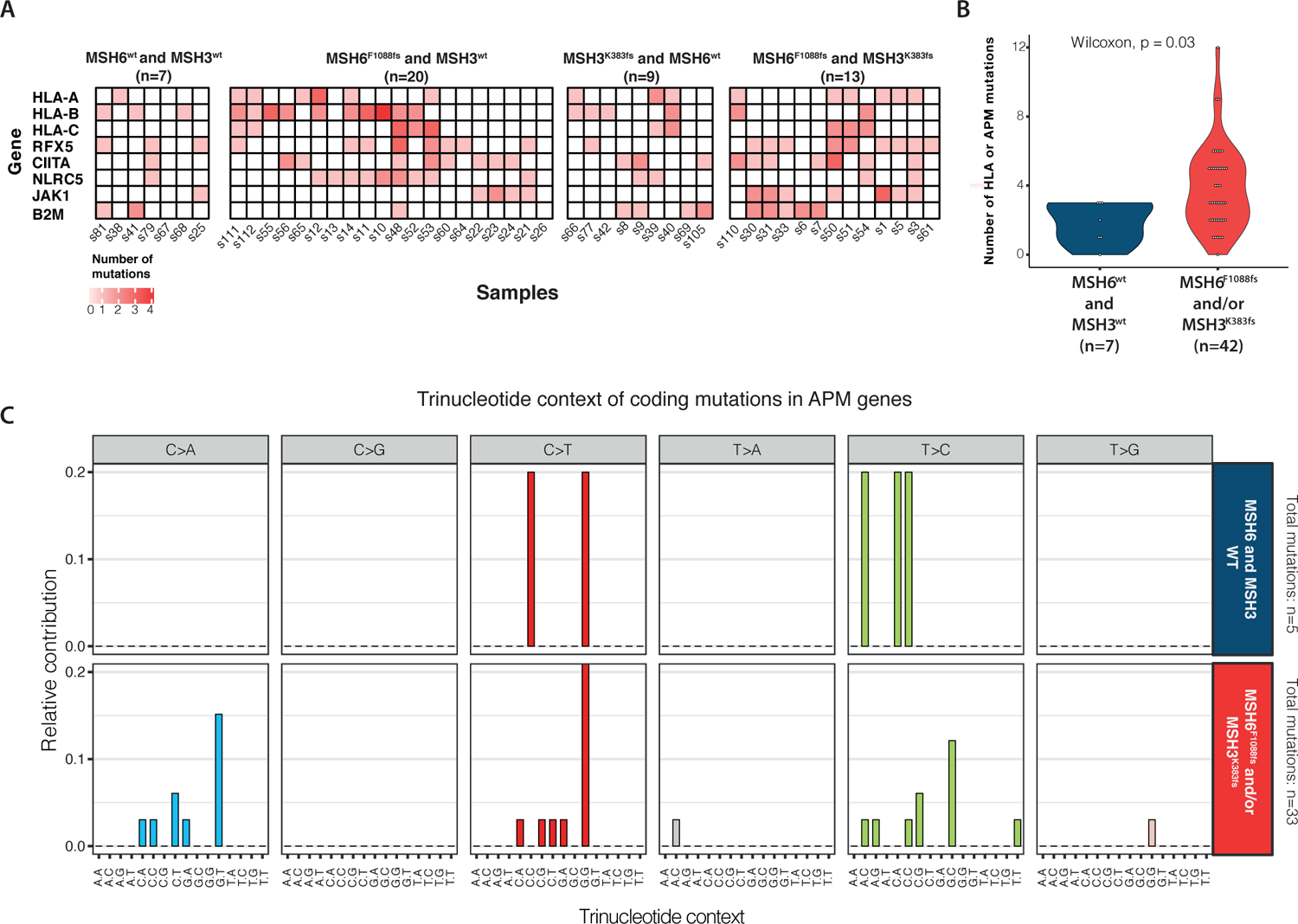
Overview antigen presentation machinery mutations. (A) Heatmaps showing number of mutations in antigen presentation machinery genes in samples according to MSH6 and MSH3 frameshift mutation status. (B) Violin plot showing increased number of HLA or antigen presentation machinery gene mutations in samples according to MSH6 and/or MSH3 homopolymer frameshifts. (C) Trinucleotide context of antigen presentation machinery gene mutations, showing increased C>A transversion contribution in samples with MSH6 and/or MSH3 homopolymer frameshift mutations.

**Figure S9.**
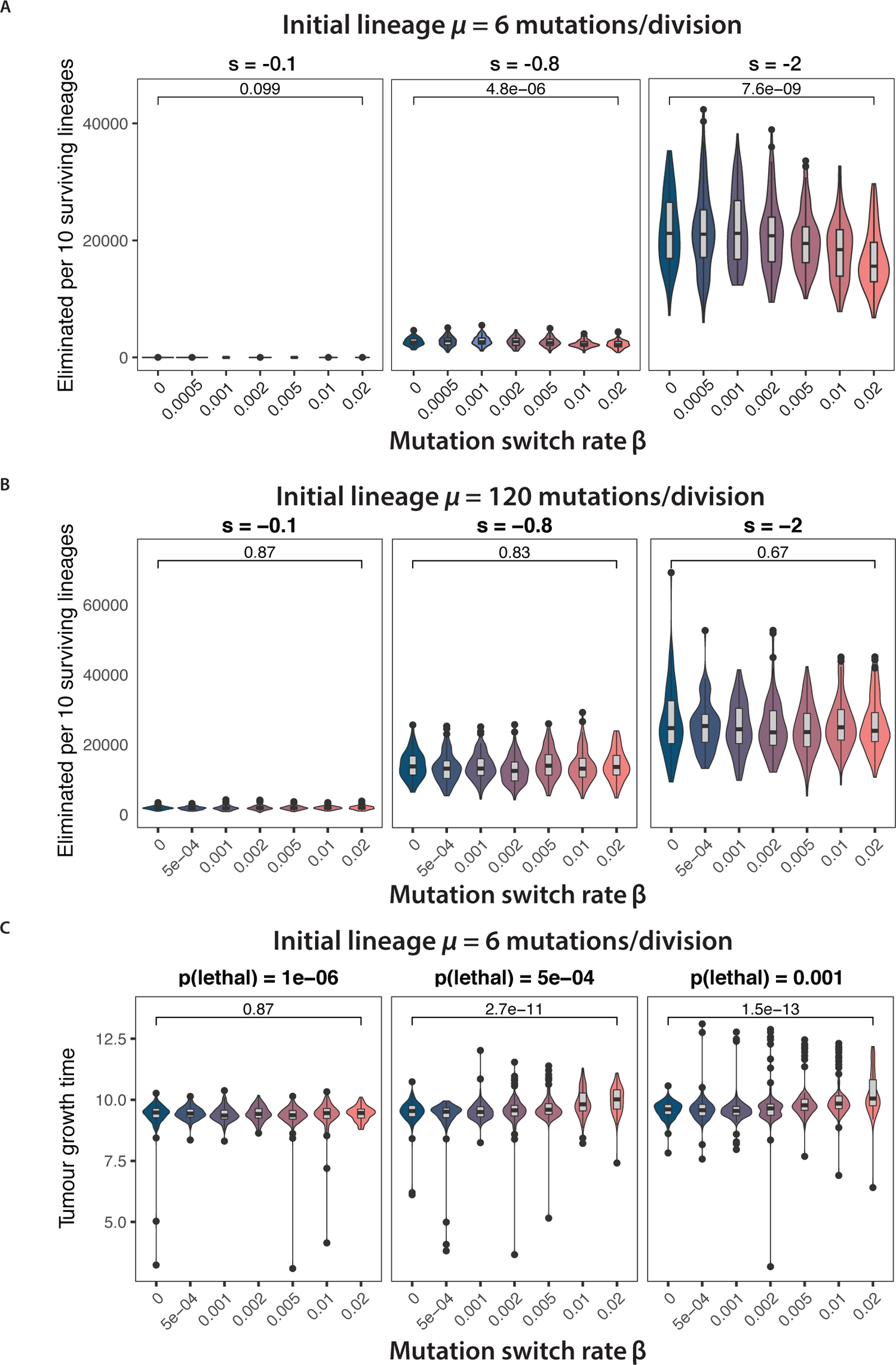
Mathematical model of stochastic mutation rate switching. (A) Number of lineages eliminated per 10 surviving lineages, computed from 100 simulated tumours at increasing immune selection (left to right) and switching rate (x-axis). Tumours are initiated with *m* = 6 mutations/division. (B) Number of lineages eliminated per 10 surviving lineages, computed from 100 simulated tumours at increasing selection rate (left to right) and mutation rate switching rate (x axis). Tumours are initiated with *m* = 120 mutations/division. (C) Tumour growth time (in arbitrary units) between establishing immune escape and reaching detectable size, computed from 100 hyper-mutated simulated tumours at increasing lethal mutation rate (left to right) and mutation rate switching rate (x axis). The Wilcoxon test statistic is reported on each panel.

**Figure S10.**
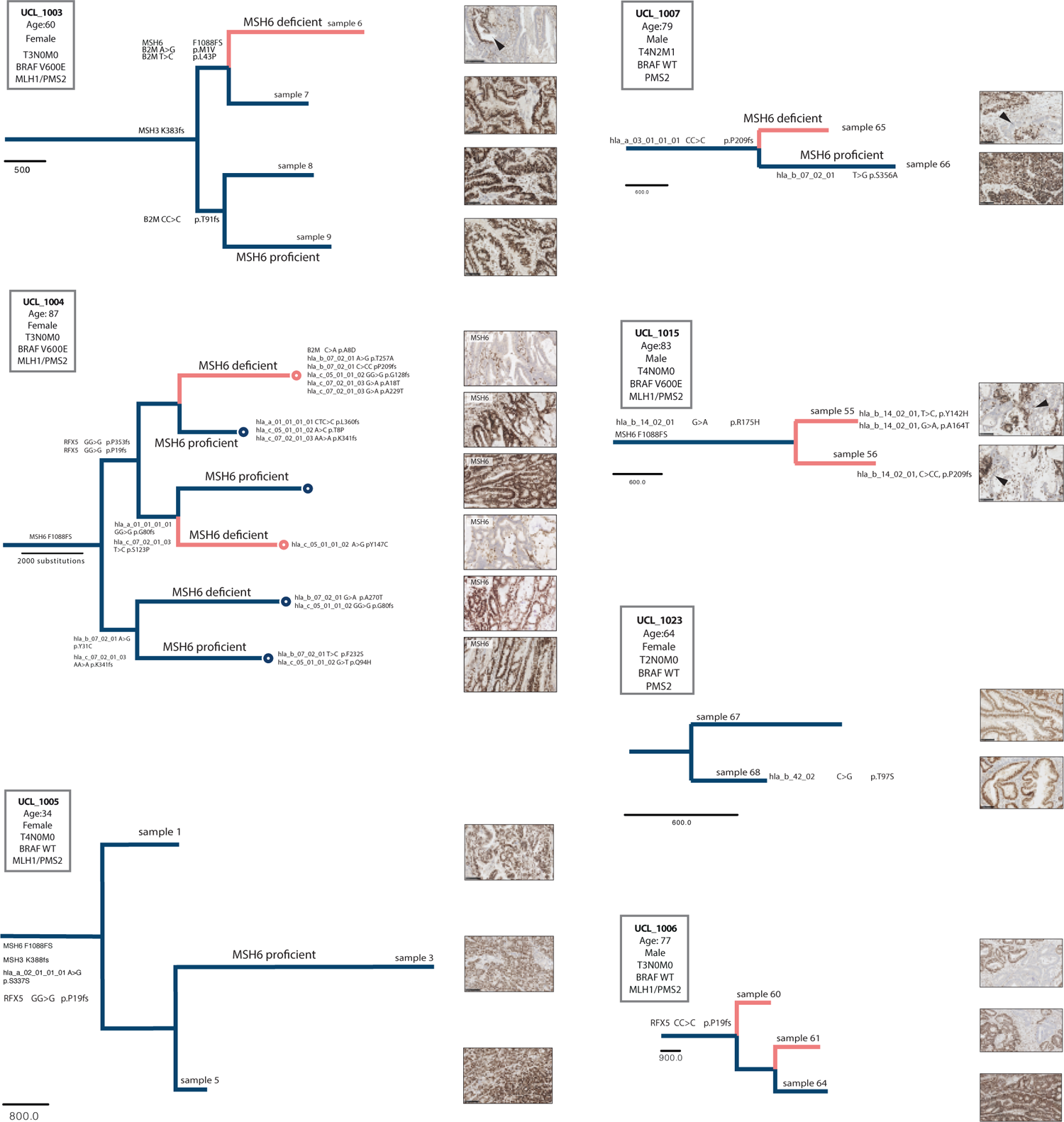
Phylogenetic trees of tumours not displayed in main figures.

**Figure S11.**
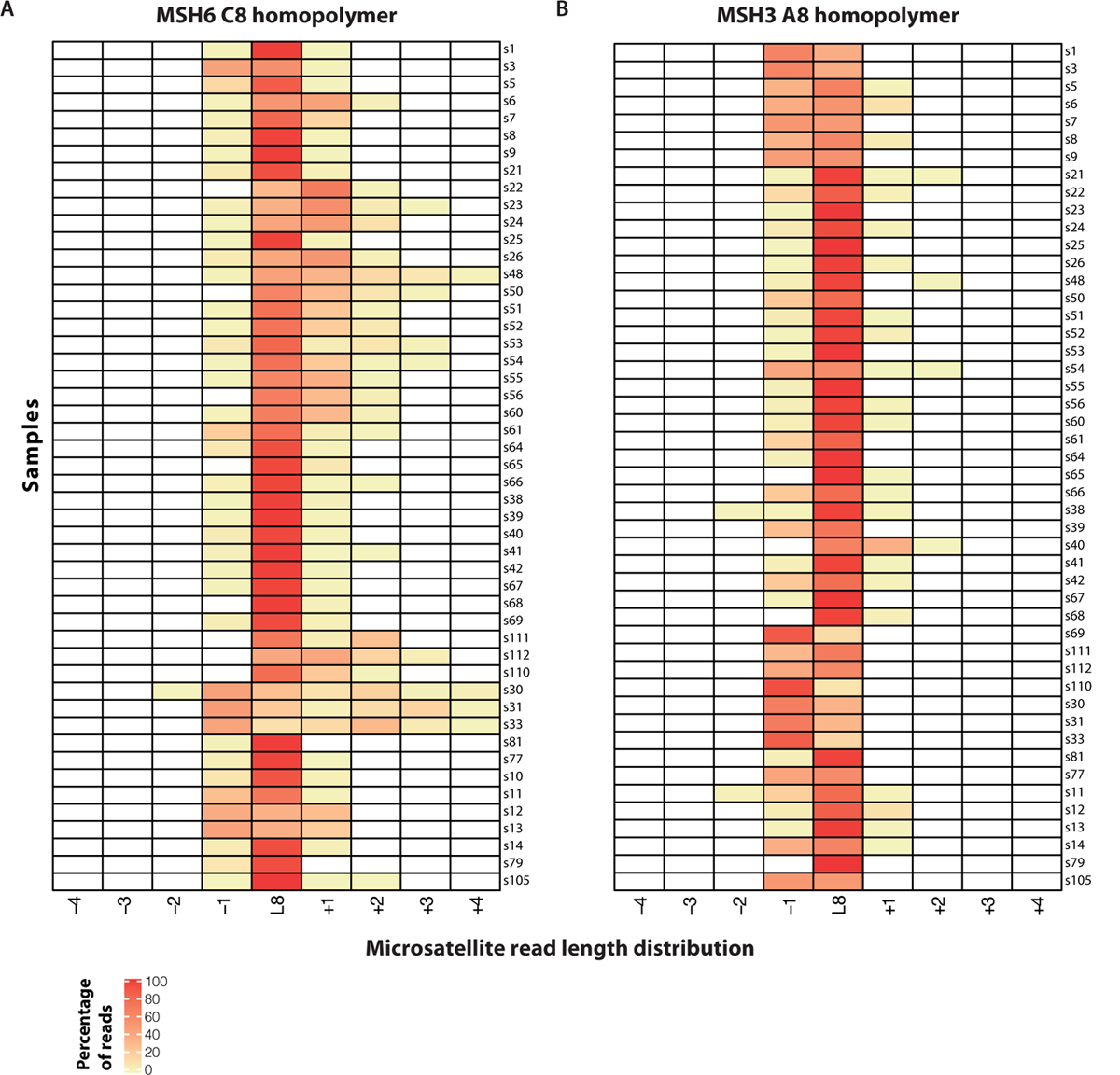
Microsatellite length distribution heatmap. (A-B) Microsatellite length distribution heatmap for the MSH6 C8 (left) and MSH3 A8 (right) homopolymers in all samples (n=49).

**Figure S12.**
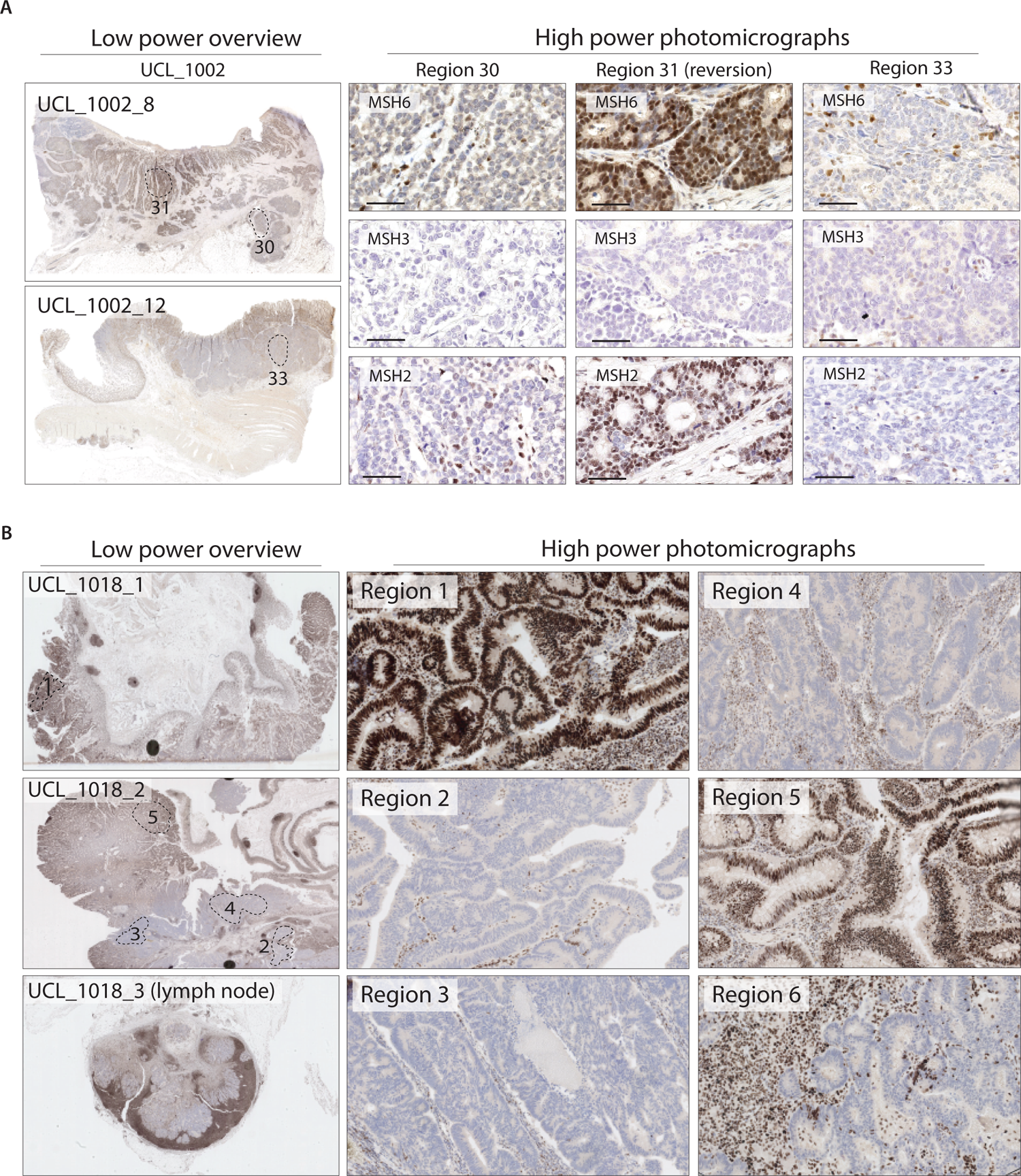
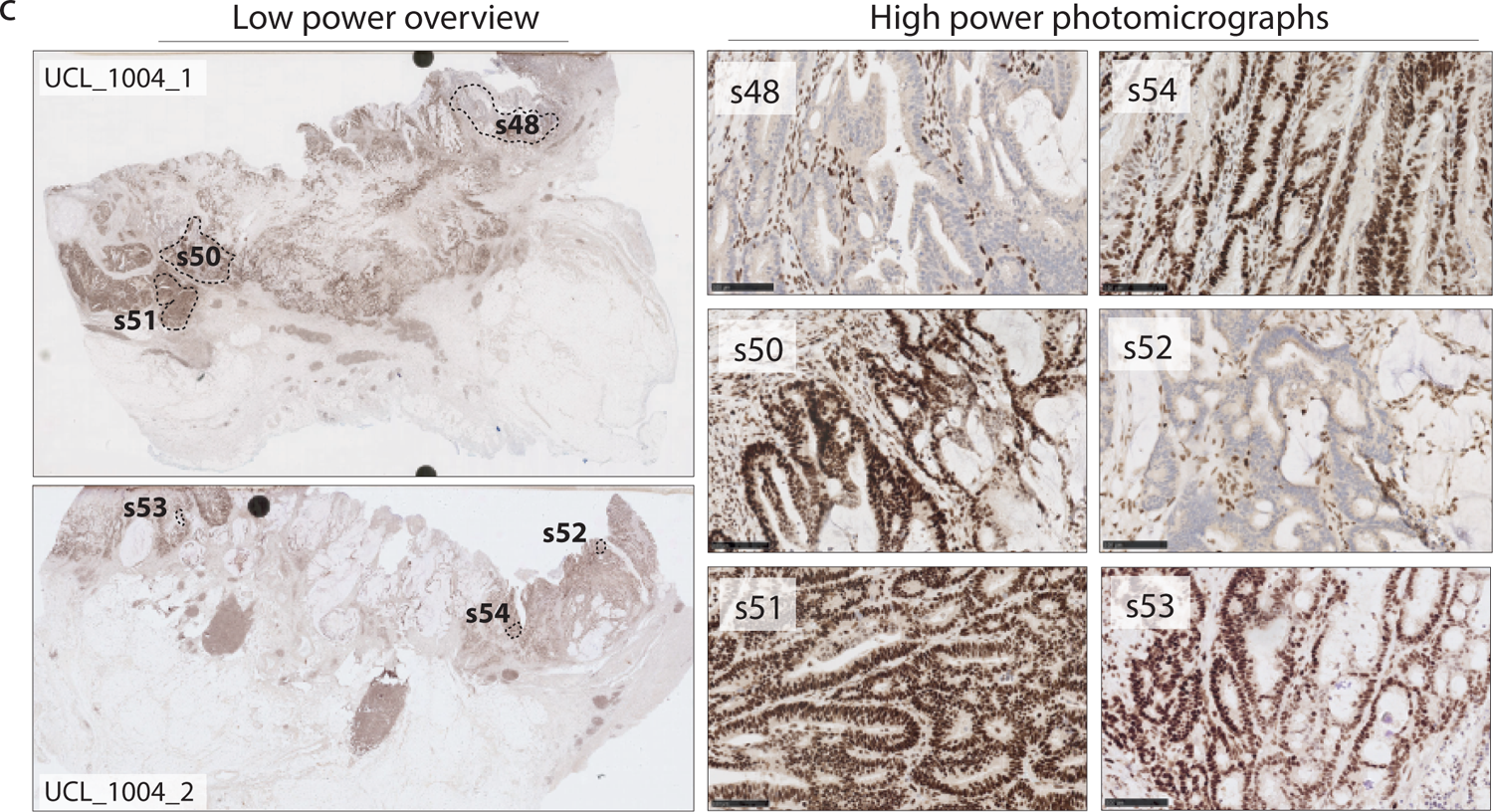
MSH6 immunohistochemistry for cases shown in main Figure 5 and Supplementary Figure S10. (A) Tumour UCL_1002 (B) Tumour UCL_1018 and (C) Tumour UCL_1004.

## Supplementary Tables

**Table S1.** Multiple linear regression analysis of association between frameshifts in MSI targets and total mutation burden (A) Model with 23 target coding microsatellites including MSH6 and MSH3 (B) Model with MSH6 and MSH3 frameshifts individually and combined.

| Independent variables | Homopolymer | Cases mutated | Beta coefficient | Standard error | t value | P value |
| --- | --- | --- | --- | --- | --- | --- |
| MSH3 | A8 | 143 (66%) | 57294 | 16962.2 | 3.378 | 0.000886 |
| MSH6 | C8 | 81 (37%) | 47190.3 | 16039.6 | 2.942 | 0.003663 |
| RFX5 | G7 | 29 (13%) | -6426.2 | 22979.3 | -0.28 | 0.780049 |
| MBD4 | T10 | 100 (46%) | -226.7 | 15291 | -0.015 | 0.988185 |
| AIM2 | T10 | 184 (85%) | 217.7 | 21976.5 | 0.01 | 0.992106 |
| ACVR2A | A8 | 212 (98%) | -60886.3 | 56695.4 | -1.074 | 0.284214 |
| DOCK3 | C7 | 128 (59%) | 18511.4 | 16435.9 | 1.126 | 0.261462 |
| TGFBR2 | A10 | 205 (94%) | 58377.2 | 37461 | 1.558 | 0.120807 |
| GLYR1 | C8 | 125 (58%) | 9028.4 | 15735.2 | 0.574 | 0.566801 |
| OR51E | A8 | 66 (30%) | -1741.2 | 16822.1 | -0.104 | 0.917668 |
| CLOCK | A9 | 140 (65%) | 14815.2 | 17093.2 | 0.867 | 0.387177 |
| CASP5 | T8 | 15 (7%) | -39338.4 | 30024.3 | -1.31 | 0.191695 |
| JAK1 | T8 | 21 (10%) | 21234.3 | 26056.1 | 0.815 | 0.416118 |
| TAF1B | A11 | 208 (96%) | 34930.8 | 42449.4 | 0.823 | 0.411602 |
| BAX | G8 | 115 (53%) | 24170.5 | 15973.8 | 1.513 | 0.131898 |
| MYH11 | G8 | 98 (45%) | -15812.1 | 15776.1 | -1.002 | 0.317476 |
| HPS1 | G8 | 82 (38%) | 21541.7 | 15546.3 | 1.386 | 0.167471 |
| SLAMF1 | T9 | 87 (40%) | 14910.3 | 15689 | 0.95 | 0.343125 |
| HNF1A | C8 | 22 (10%) | 4553.4 | 26119.2 | 0.174 | 0.861788 |
| RGS12 | A9 | 72 (33%) | 12400.9 | 16625.4 | 0.746 | 0.456646 |
| ELAVL3 | C9 | 85 (39%) | -6014.3 | 15507.7 | -0.388 | 0.698578 |
| SMAP1 | A10 | 183 (84%) | 38820.9 | 22473.8 | 1.727 | 0.085715 |
| SLC22A9 | A11 | 206 (95%) | 62795.6 | 37207.8 | 1.688 | 0.093101 |
| Tumour purity | - | - | 178452.1 | 58599.6 | 3.045 | 0.002653 |
| Patient age | - | - | 792.4 | 608.5 | 1.302 | 0.1944 |
|  |  |  | **R<sup>2</sup> (adjusted)** |  |  | 0.225 |
|  |  |  | **P value** |  |  | 4.20x10<sup>-7</sup> |

**Table S1. Multiple linear regression analysis of association between frameshifts in MSI targets and total mutation burden | (A)** Model with 23 target coding homopolymers including MSH6 and MSH3 (B) Model with MSH6 and MSH3 frameshifts individually and combined.
| Independent variables | Beta coefficient | Standard error | t value | P value |
| --- | --- | --- | --- | --- |
| MSH3 K383fs | 88038.2 | 20078.9 | 4.385 | 0.0000183 |
| MSH6 F1088fs | 63675.6 | 27247.8 | 2.337 | 0.0204 |
| MSH6 F1088fs & MSH3 K383fs | 139338.8 | 22163.6 | 6.287 | 1.83x10<sup>-9</sup> |
| tumour purity | 145461.6 | 56173.8 | 2.589 | 0.0103 |
| patient age | 874 | 591.4 | 1.478 | 0.141 |
| **R<sup>2</sup> (adjusted)** |  |  |  | 0.17 |
| **p-value** |  |  |  | 1.87x10<sup>-8</sup> |

**Table S2.**
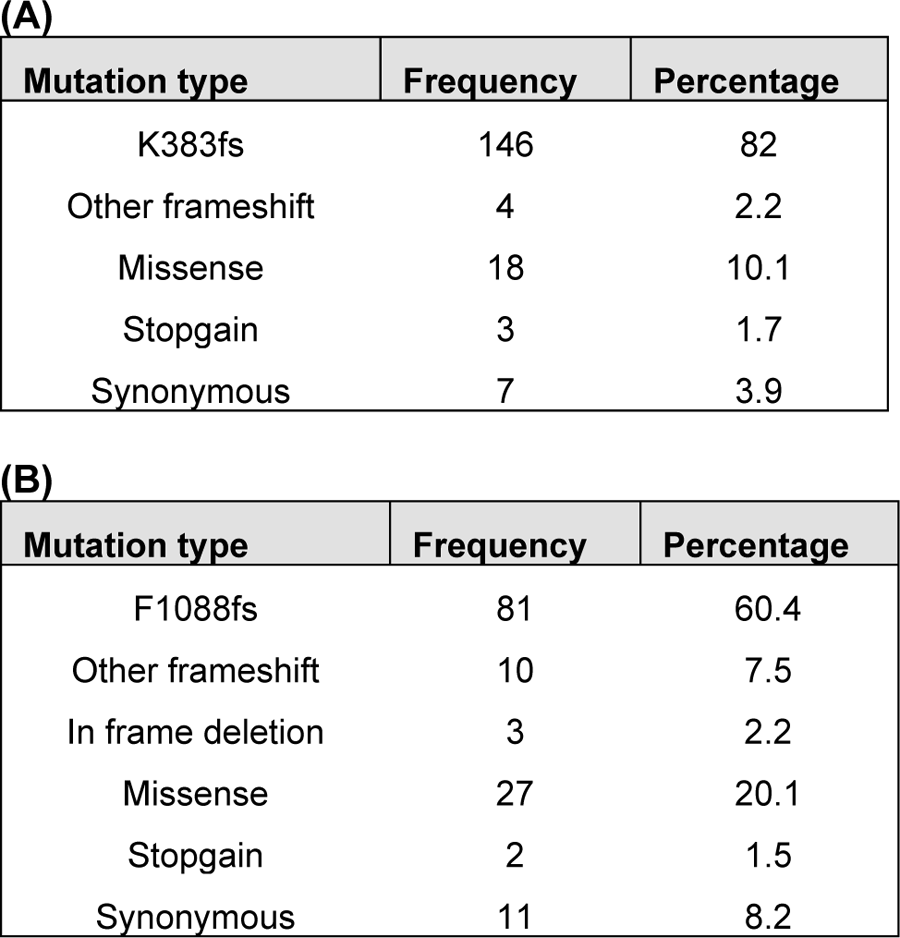
Frequency of all coding mutations in GEL CRC MSI cohort for (A) MSH3 and (B) MSH6

**(A)**
| **Mutation type** | **Frequency** | **Percentage** |
| --- | --- | --- |
| K383fs | 146 | 82 |
| Other frameshift | 4 | 2.2 |
| Missense | 18 | 10.1 |
| Stopgain | 3 | 1.7 |
| Synonymous | 7 | 3.9 |

| **Mutation type** | **Frequency** | **Percentage** |
| --- | --- | --- |
| F1088fs | 81 | 60.4 |
| Other frameshift | 10 | 7.5 |
| In frame deletion | 3 | 2.2 |
| Missense | 27 | 20.1 |
| Stopgain | 2 | 1.5 |
| Synonymous | 11 | 8.2 |

**Table S3.**
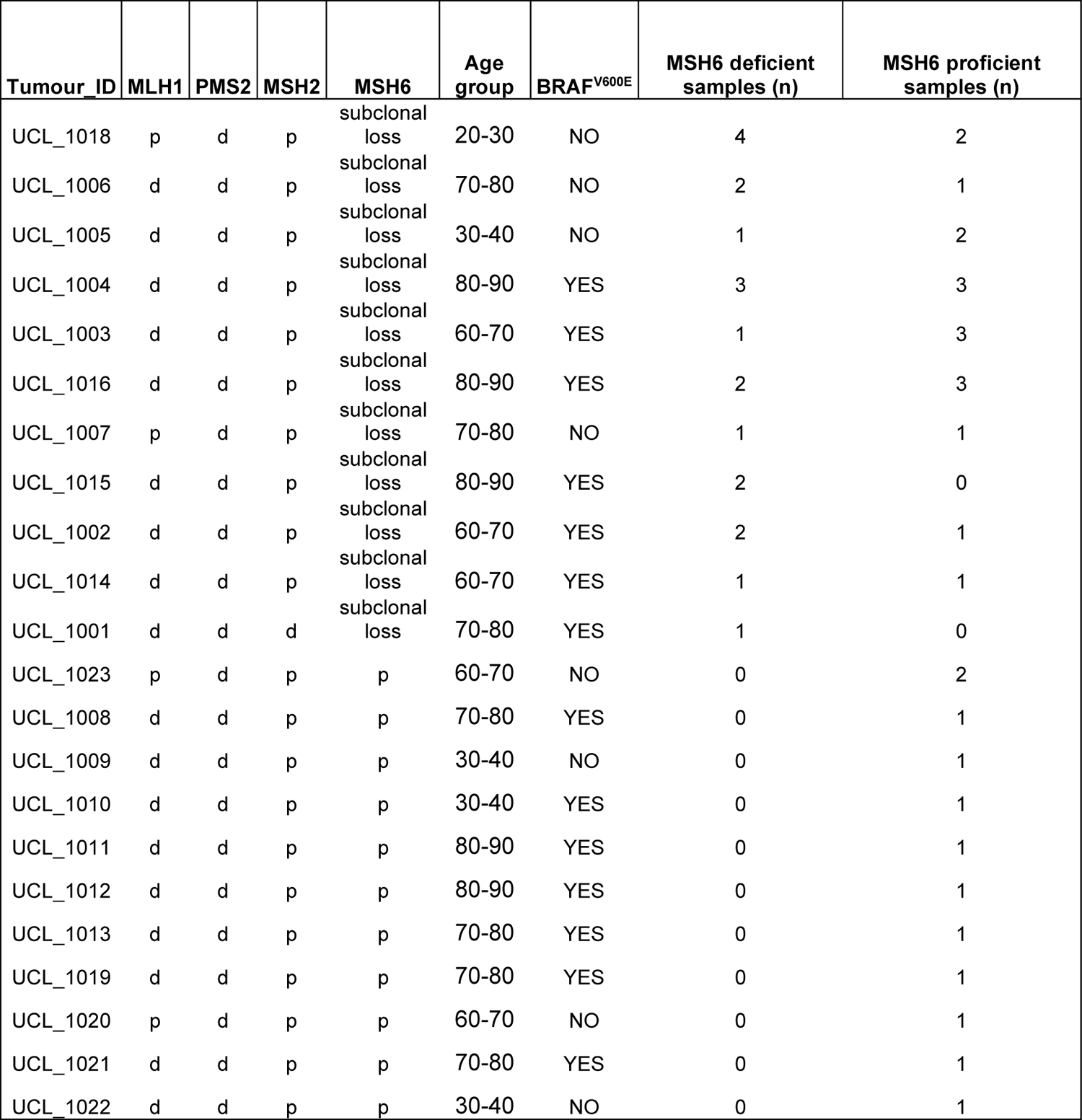
Clinical and pathological characteristics of 22 tumours used in whole exome sequencing experiments.

**Table S4.** Linear mixed effect modelling of tumour mutation burden

| AIC | BIC | logLik | deviance | df.resid |
| --- | --- | --- | --- | --- |
| 881.9 | 893.2 | -434.9 | 869.9 | 43 |

Random effects:
| Groups | Name | Std.Dev. |
| --- | --- | --- |
| Tumours | (Intercept) | 2209 |
| Residual |  | 1061 |

Number of obs: 49, groups: Tumours, 22 Fixed Effects:
| (Intercept) | MSH6_MSH3_status | purity | age |
| --- | --- | --- | --- |
| 1509.53 | 612.23 | 1580.30 | 29.67 |

| AIC | BIC | logLik | deviance | df.resid |
| --- | --- | --- | --- | --- |
| 884.5813 | 894.0404 | 437.2906 | 874.5813 | 44 |

Random effects:
| Groups | Name | Std.Dev. |
| --- | --- | --- |
| Tumours | (Intercept) | 2428 |
| Residual |  | 1080 |

Number of obs: 49, groups: Tumours, 22 Fixed Effects:
| (Intercept) | MSH6_MSH3_status | purity | age |
| --- | --- | --- | --- |
|  | 3065.10 | 29.91 | 1491.92 |

|  | npars | AIC | BIC | logLik | deviance | Chisq | Df | Pr(>Chisq) |
| --- | --- | --- | --- | --- | --- | --- | --- | --- |
| null_model | 5 | 884.58 | 894.04 | -437.29 | 874.58 |  |  |  |
| full_model | 6 | 881.85 | 893.20 | -434.93 | 869.85 | 4.7309 | 1 | 0.02963 * |

**Table S4. Linear mixed effect modelling of tumour mutation burden**
|  | npars | AIC | LRT | Pr(Chi) |
| --- | --- | --- | --- | --- |
| <none> |  | 881.85 |  |  |
| MSH6_MSH3_status | 1 | 884.58 | 4.7309 | 0.02963 * |
| purity | 1 | 880.80 | 0.9470 | 0.33048 |
| age | 1 | 881.17 | 1.3232 | 0.25001 |

**Table S5.** Antibody conditions and details (A) For immunohistochemistry (B) For multiplex immunofluorescence.

| Epitope | Clone | Company | Dilution | Antigen retrieval (mins) | Primary incubation (mins) | Post-primary (mins) | HRP polymer (mins) | DAB (mins) |
| --- | --- | --- | --- | --- | --- | --- | --- | --- |
| MSH6 | EP49 | Agilent | 1:400 | ER2 40 | 30 | 20 | 20 | 5 |
| MLH1 | ES05 | Agilent | 1:200 | ER2 40 | 30 | 20 | 20 | 5 |
| PMS2 | A16-4 | BD Biosciences | 1:300 | ER2 40 | 15 | 8 | 8 | 5 |
| MSH2 | FE11 | Agilent | 1:100 | ER2 40 | 30 | 20 | 20 | 5 |
| MSH3 | 611390 | BD Biosciences | 1:100 | ER1 30 | 30 | 8 | 8 | 5 |

| Opal fluorophore | Opal fluorophore dilution | Primary epitope | Clone | Company | Primary dilution | Primary incubation (mins) | Antigen retrieval |
| --- | --- | --- | --- | --- | --- | --- | --- |
| 520 | 1:150 | MSH6 | EP49 | Agilent | 1:100 | 60 | ER2 (40) |
| 540 | 1:150 | CD20 | L26 | Agilent | 1:150 | 15 | ER1(20) |
| 570 | 1:150 | FOXP3 | D608 R | Cell Signalling Technology | 1:200 | 30 | ER2(20) |
| 620 | 1:150 | CD4 | 4B12 | Agilent | 1:50 | 30 | ER1(20) |
| 650 | 1:150 | PANCK | AE1/3 | Agilent | 1:700 | 15 | ER1(20) |
| 690 | 1:150 | CD8 | 4B11 | Agilent | 1:100 | 15 | ER1(20) |

**Table S6.** Comparison of ORION cell segmentation with other methods (A) In dataset A (B) In dataset B.

| Methods | JSC | MAD | Hausdorff | DiceFP | DiceFN |
| --- | --- | --- | --- | --- | --- |
| Otsu [15] | 83.5 | 4.5 | 11.5 | **3** | 16.7 |
| Three-step [16] | 88.4 | 4.7 | 13.4 | 5.3 | 5.2 |
| LSBR [17] | 83.2 | 5.8 | 19.8 | 11.8 | 9.1 |
| LLBWIP [18] | 91.6 | 3.5 | 12.7 | 4.7 | 3.9 |
| RFOVE [7] | 89.8 | **2.8** | 7.5 | 5.2 | 5.7 |
| **ORION (proposed)** | **92.4** | 3.6 | **6.8** | 5.4 | **3.2** |

| Methods | JSC | MAD | Hausdorff | DiceFP | DiceFN |
| --- | --- | --- | --- | --- | --- |
| Otsu [15] | 81.4 | 5.8 | 17.5 | **3.9** | 17.1 |
| Watershed [2] | 85.1 | 4.9 | 10.1 | 4.9 | 10.3 |
| Mask R-CNN [12] | 83.7 | 5.1 | 10.7 | 5.2 | 11.8 |
| **ORION (proposed)** | **88.2** | **4.2** | **6.2** | 4.5 | **8.8** |

**Table S7.**
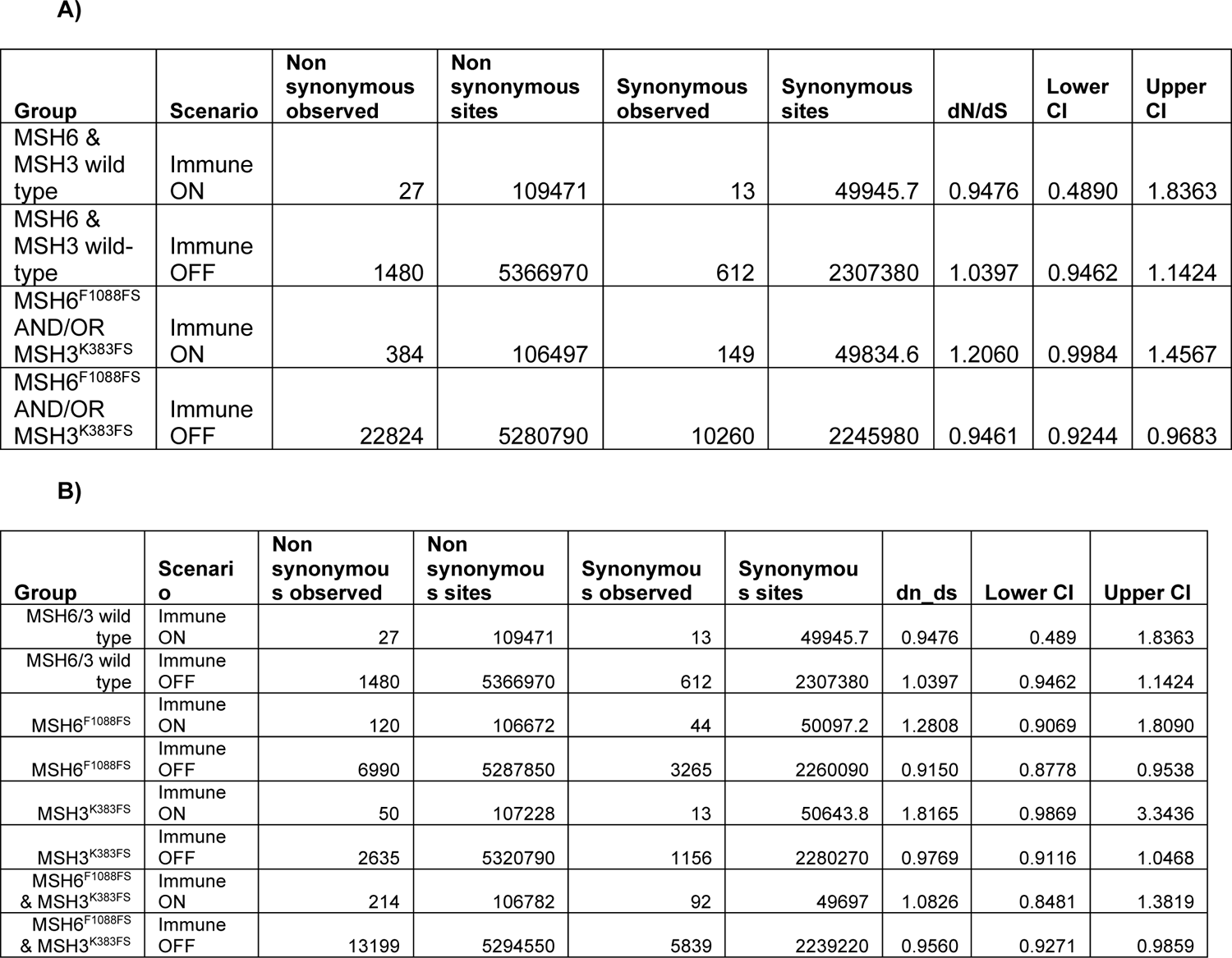
Data used for calculation of immune ON and OFF dN/dS values A) Samples grouped into two categories according to MSH6 ^F1088FS^ /MSH3 ^K383FS^ status as presented in figure 6A B) Samples grouped into 4 categories according to presence of MSH6 ^F1088FS^, MSH3 ^K383FS^ or both MSH6 ^F1088FS^/MSH3 ^K383FS^ status.

| Group | Scenario | Non synonymous observed | Non synonymous sites | Synonymous observed | Synonymous sites | dN/dS | Lower CI | Upper CI |
| --- | --- | --- | --- | --- | --- | --- | --- | --- |
| MSH6 & MSH3 wild type | Immune ON | 27 | 109471 | 13 | 49945.7 | 0.9476 | 0.4890 | 1.8363 |
| MSH6 & MSH3 wild-type | Immune OFF | 1480 | 5366970 | 612 | 2307380 | 1.0397 | 0.9462 | 1.1424 |
| MSH6<sup>F1088FS</sup> AND/OR MSH3<sup>K383FS</sup> | Immune ON | 384 | 106497 | 149 | 49834.6 | 1.2060 | 0.9984 | 1.4567 |
| MSH6<sup>F1088FS</sup> AND/OR MSH3<sup>K383FS</sup> | Immune OFF | 22824 | 5280790 | 10260 | 2245980 | 0.9461 | 0.9244 | 0.9683 |

**Table S7.** Data used for calculation of immune ON and OFF dn/ds values A) Samples grouped into two categories according to MSH6<sup>F1088FS</sup> /MSH3<sup>K383FS</sup> status as presented in figure 6A B) Samples grouped into 4 categories according to presence of MSH6<sup>F1088FS</sup>, ...
| Group | Scenario | Non synonymous observed | Non synonymous sites | Synonymous observed | Synonymous sites | dn_ds | Lower CI | Upper CI |
| --- | --- | --- | --- | --- | --- | --- | --- | --- |
| MSH6/3 wild type | Immune ON | 27 | 109471 | 13 | 49945.7 | 0.9476 | 0.489 | 1.8363 |
| MSH6/3 wild type | Immune OFF | 1480 | 5366970 | 612 | 2307380 | 1.0397 | 0.9462 | 1.1424 |
| MSH6<sup>F1088FS</sup> | Immune ON | 120 | 106672 | 44 | 50097.2 | 1.2808 | 0.9069 | 1.8090 |
| MSH6<sup>F1088FS</sup> | Immune OFF | 6990 | 5287850 | 3265 | 2260090 | 0.9150 | 0.8778 | 0.9538 |
| MSH3<sup>K383FS</sup> | Immune ON | 50 | 107228 | 13 | 50643.8 | 1.8165 | 0.9869 | 3.3436 |
| MSH3<sup>K383FS</sup> | Immune OFF | 2635 | 5320790 | 1156 | 2280270 | 0.9769 | 0.9116 | 1.0468 |
| MSH6<sup>F1088FS</sup> & MSH3<sup>K383FS</sup> | Immune ON | 214 | 106782 | 92 | 49697 | 1.0826 | 0.8481 | 1.3819 |
| MSH6<sup>F1088FS</sup> & MSH3<sup>K383FS</sup> | Immune OFF | 13199 | 5294550 | 5839 | 2239220 | 0.9560 | 0.9271 | 0.9859 |

